# A Hotspot Phosphorylation Site on SHP2 Drives Oncoprotein Activation and Drug Resistance

**DOI:** 10.1101/2025.06.11.659120

**Authors:** Prashath Karunaraj, Remkes Scheele, Malcolm L. Wells, Ruchita Rathod, Sophia Abrahamson, Lila C. Taylor, Ipek S. Gokulu, Lamia Chowdhury, Abiha Kazmi, Weixiao Song, Peter Hornbeck, Jing Li, Anum Glasgow, Neil Vasan

## Abstract

SHP2 is a phosphatase and a critical mediator of receptor tyrosine kinase (RTK)-driven RAS/mitogen-activated protein kinase (MAPK) signaling. Despite promising preclinical data, SHP2 inhibitors have shown minimal clinical efficacy, with no defined clinical mechanisms of primary resistance. Here, we elucidate phosphorylation of SHP2 at tyrosine 62 (pY62) as a hotspot phosphorylation site in the proteome and RTK-driven tumor types in patients. We demonstrate that SRC family kinases directly phosphorylate SHP2 at Y62, downstream of but not directly phosphorylated by RTKs. Using biochemical and biophysical analyses, we show that SHP2 Y62D enforces an open, active conformation, resulting in constitutive phosphatase activation that is sufficient to activate MAPK signaling and confer resistance to allosteric SHP2 inhibitors. These findings establish that SHP2 pY62 is a phosphorylation hotspot phenocopying mutational activation, a mechanism of primary resistance to SHP2 inhibitors, and a cancer drug target distinct from wildtype SHP2.

**Statement of significance:** This study identifies phosphorylation of SHP2 at tyrosine 62 (pY62) as a conserved mechanism of resistance to allosteric SHP2 inhibitors. By stabilizing an open, active SHP2 conformation, pY62 phenocopies oncogenic *PTPN11* mutations and sustains MAPK signaling across cancer types. These findings redefine SHP2 inhibitor resistance as a phosphorylation-driven, target-intrinsic process, nominate pY62 as a potential biomarker for therapeutic response, and propose phosphorylated SHP2 as a distinct drug target.

## Introduction

SHP2, encoded by the *PTPN11* gene, is a non-receptor protein tyrosine phosphatase^1–5^ that serves as a critical mediator of receptor tyrosine kinase (RTK) signaling to the RAS/MAPK pathway^6,7^. In its basal state, SHP2 adopts an autoinhibited closed conformation, in which the N-terminal Src homology 2 (SH2) domain occludes the protein tyrosine phosphatase (PTP) domain active site^8,9^ (**Fig. 1A**). Upon activation of RTKs, SHP2 binds to phosphotyrosine-containing motifs through its SH2 domains, relieving autoinhibition and enabling its catalytic activity. In addition to its phosphatase function, SHP2 acts as a signaling scaffold, integrating RTK inputs and facilitating the activation of downstream effectors such as RAS and ERK^10–15^.

**Figure 1:**
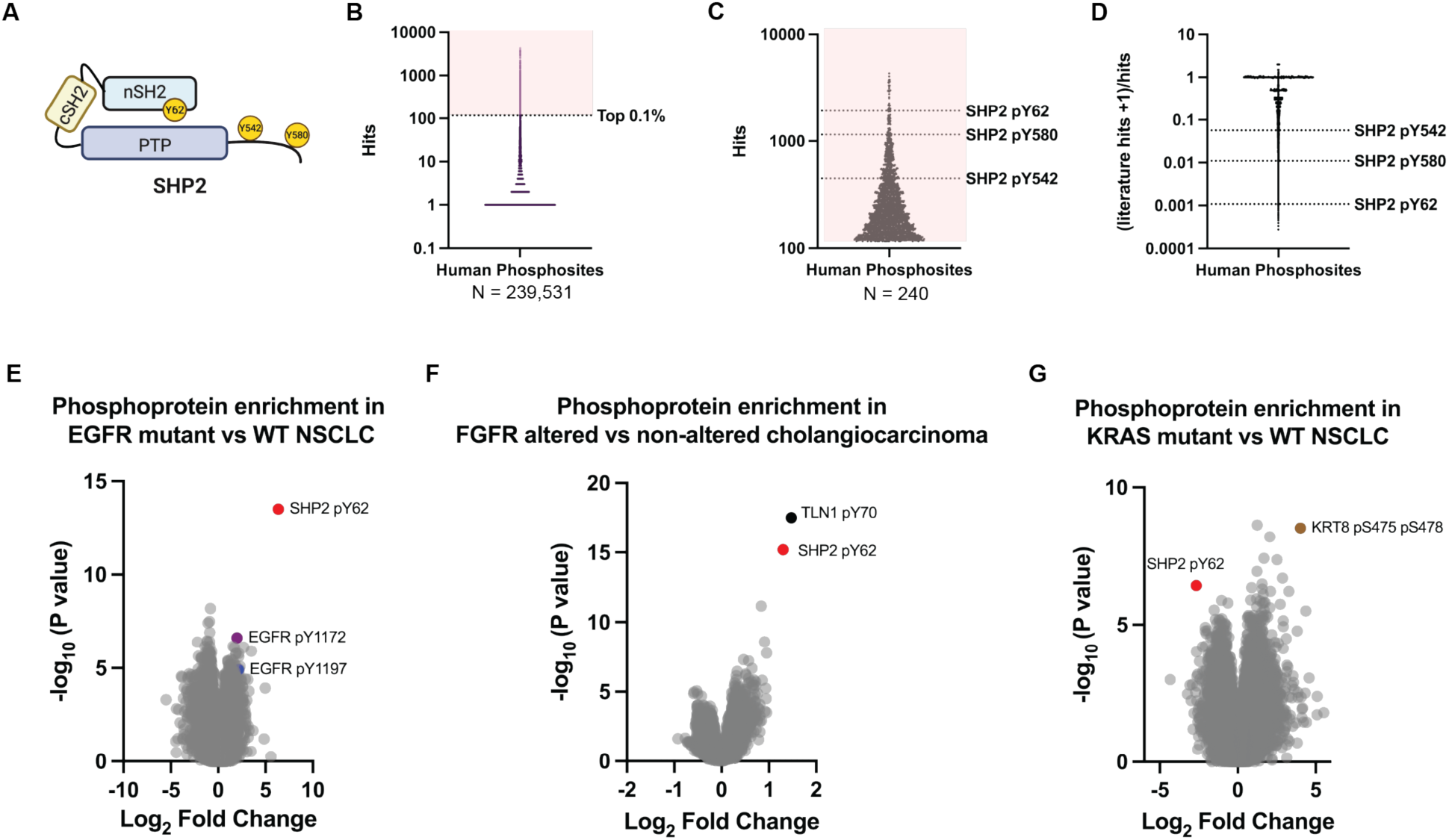
SHP2 Y62 is one of the most abundantly phosphorylated human proteins and is enriched in diverse RTK-driven tumor types. (**A)** Domain schematic of SHP2 protein with phosphosites indicated. (**B)** Human phosphosite frequency (PhosphoSitePlus) with (**C**) top 0.1% (n=240) hits shown and SHP2 phosphosites indicated. (**D)** Published literature frequency of SHP2 phosphosites. (**E**-**G**) Volcano plots of phosphoprotein abundances in RTK-altered human tumors compared to wildtype counterparts (CPTAC) in (**E**) EGFR-mutated non-small cell lung cancer, (**F**) FGFR-altered cholangiocarcinoma, and (**G**) KRAS-mutated non-small cell lung cancer.

Activating mutations in SHP2 have been well characterized as oncogenic drivers in hematologic malignancies, including juvenile myelomonocytic leukemia (JMML) and subsets of acute myeloid leukemia (AML), and as germline mutations which underlie Noonan syndrome (NS)^16^. More recently, SHP2 has emerged as an essential signaling node in RTK-driven solid tumors, where SHP2 is rarely mutated^17–19^. Functional genetic studies have demonstrated that SHP2 is required for survival in RTK-driven cancer models, including those driven by oncogenic epidermal growth factor receptor (EGFR), anaplastic lymphoma kinase (ALK), and fibroblast growth factor receptor (FGFR) alterations^20^. These findings catalyzed the development of allosteric SHP2 inhibitors (SHP2i) that function as molecular glues, stabilizing the autoinhibited conformation of the protein (e.g., SHP099, RMC4550, TNO155)^21^. Although these agents have demonstrated potent preclinical activity, clinical trials to date have reported negligible objective response rates (ORR = ∼0%)^22–24^ with stable disease at best, and have been associated with dose-limiting toxicities, including fatigue, rash, and cytopenias. These challenges highlight the need to investigate mechanisms of primary resistance to SHP2i and identify biomarkers that modulate SHP2 dependence and therapeutic response.

Considerable preclinical effort has gone into uncovering genetic resistance mechanisms to SHP2i including activating SHP2 mutations that lock the protein in an open, active state, making it resistant to allosteric inhibitors designed to bind the inactive form. However, activating mutations are rare in RTK-driven solid tumors. Additionally, much attention has focused on phosphorylation of SHP2 at its canonical C-terminal tyrosines, Y542 and Y580, which are known to promote scaffolding interactions and downstream signaling^15,25–27^, but their direct effect on SHP2’s catalytic activity remains unclear^28^.

Despite the clinical momentum behind SHP2 inhibitors, emerging evidence points to alternative phosphorylation sites, including SHP2 Y62 (pY62) as potential modulators of inhibitor sensitivity. While pY62 has been investigated in select contexts, including cytokine-stimulated hematopoietic stem cells and drug-resistant leukemia cells^29–31^, where it may stabilize the open, active conformation and confers resistance to the allosteric inhibitor SHP099^30^, its broader relevance remains unknown. Specifically, the prevalence of SHP2 pY62 across diverse tumor types, its cellular regulation, and its biochemical and structural impact on SHP2 phosphatase function remain unknown or incomplete. Our interest in SHP2 pY62 was driven by two independent observations that underscore its potential significance. First, a comprehensive phosphoproteomic analysis identified SHP2 pY62 as one of the most recurrent phosphorylation events in the human proteome. Second, SHP2 pY62 is one of the most differentially enriched phosphopeptides across multiple RTK-driven tumor types in patient samples. This set of observations in normal cells and tissues, cancer cells, and patient samples prompted us to investigate the functional, mechanistic, and therapeutic consequences of SHP2 Y62 phosphorylation.

## Results

### SHP2 pY62 is a hotspot phosphorylation in normal cells/tissues and cancer cells and is enriched in diverse RTK-driven tumor types

Inspired by cancer genomics efforts to define frequent “mutational hotspots”^32–34^, we performed a comprehensive analysis of the human phosphoproteome using PhosphositePlus^31^. We identified 239,531 human protein phosphorylation sites (phosphosites) across normal cells and tissues, and cancer cells (**Fig. 1B**) (**Supp. Data 1**). We posited that the most frequent phosphosites may be enriched for functionality, and focused analysis on the top 240 (0.1%) of phosphosites. In line with this assertion, many of the most frequent phosphosites were known kinase activation loop sites including GSK3α pY279, GSK3β pY216, and ERK2 pY187 (**Supp. Data 1**). We examined proteins with multiple phosphorylation sites and found that the top differentially phosphorylated protein was not a kinase, but the protein phosphatase SHP2 (**Fig. 1C**). The known SHP2 C-terminal tyrosine phosphosites Y542 and Y580 (pY542, pY580) (**Fig. 1A**) were abundant, but surprisingly, the most abundant SHP2 phosphosite was the tyrosine Y62 (pY62) (**Fig. 1C**).

In a survey of published literature, SHP2 pY62 was reported far less frequently than the canonical C-terminal phosphorylation sites pY542 and pY580, as reflected by a markedly lower ratio of literature mentions to curated PhosphoSite records (**Fig. 1D**). Despite this underrepresentation, SHP2 Y62 is evolutionarily conserved across vertebrates, down to *Xenopus* (**Supp. Fig. 1**). Notably, SHP2 Y62 is not conserved in the closely related human phosphatase SHP1, where it aligns with phenylalanine 60 (F60), a non-phosphorylatable residue. In contrast, the adjacent SHP2 Y63 is conserved and aligns with SHP1 Y61, indicating selective evolutionary retention of the neighboring tyrosine, rather than the Y62-equivalent site. SHP2 Y62 is located in a structured core on a loop between the βD and βE strands of the SHP2 N-terminal SH2 (nSH2) domain^8^. Among the 110 SH2-domain containing proteins in humans, Y62-homologous tyrosines are present in 12 proteins, some of which are frequently phosphorylated (**Supp. Fig. 1**)^35^.

Protein phosphorylation can differ between cancer cell lines and tumors^36^. To understand the role of SHP2 pY62 in patients, we determined its abundance and enrichment across genomically defined human tumor types based on publicly available data from the Clinical Proteomic Tumor Analysis Consortium (CPTAC)^37^. SHP2 pY62 has been reported as the single most differentially increased phosphosite in patients with EGFR-mutated non-small cell lung cancer (NSCLC), compared to their wild-type (WT) counterparts^38^ (**Fig. 1E**). In fact, SHP2 pY62 was more increased than phosphorylated EGFR itself (**Fig. 1E**, **Supp. Data 2**). Additionally, SHP2 pY62 was the second most differentially increased phosphosite in patients with FGFR2-altered cholangiocarcinoma^39^ (**Fig. 1F**), and differentially increased in CPTAC datasets in patients with EGFR-amplified head and neck squamous cell carcinoma^40^, EGFRvIII glioblastoma^41^, and EGFR-altered high grade glioma^42^, compared to WT counterparts. In contrast, SHP2 pY62 was one of the most significantly decreased phosphosites in KRAS-mutant NSCLC relative to wild-type NSCLC^38^ (**Fig. 1G**, **Supp. Data 2**). Our identification of SHP2 pY62 as one of the most frequently detected phosphosites in the human phosphoproteome, along with its evolutionary conservation and selective retention across orthologs and SH2 domains, and its high frequency in RTK-driven patient tumor types suggests that Y62 phosphorylation may exert distinct conformational or functional effects that contribute to RTK-driven oncogenesis.

### SHP2 pY62 is downstream of RTKs, but Y62 is not phosphorylated by RTKs

Given the high abundance of pY62 across RTK-driven tumors, we hypothesized that RTKs signal to and directly phosphorylate SHP2 Y62 in cells (**Fig. 2A**). To test this hypothesis, we developed a phosphospecific SHP2 pY62 antibody and validated it in SHP2 knock out (KO) cells overexpressing SHP2 and the phosphodead mutant Y62F (**Supp. Fig. 2A, B**). We serum-starved U-2 OS (osteosarcoma) and BT20 (EGFR-amplified triple-negative breast cancer) cells and exposed them to various RTK ligands (EGF, FGF1, PDGF). RTK stimulation led to RTK phosphorylation at 5-10 minutes, with eventual detection of SHP2 pY62 much later, suggesting SHP2 Y62 phosphorylation is downstream of RTKs (**Fig. 2B**, **Supp. Fig. 2C-E**). However, when we exposed unstimulated cells to RTK inhibitors against EGFR, FGFR, and PDGFR in normal media, there was no decrease in SHP2 pY62 (**Fig. 2C**), showing that SHP2 Y62 is not directly phosphorylated by RTKs.

**Figure 2:**
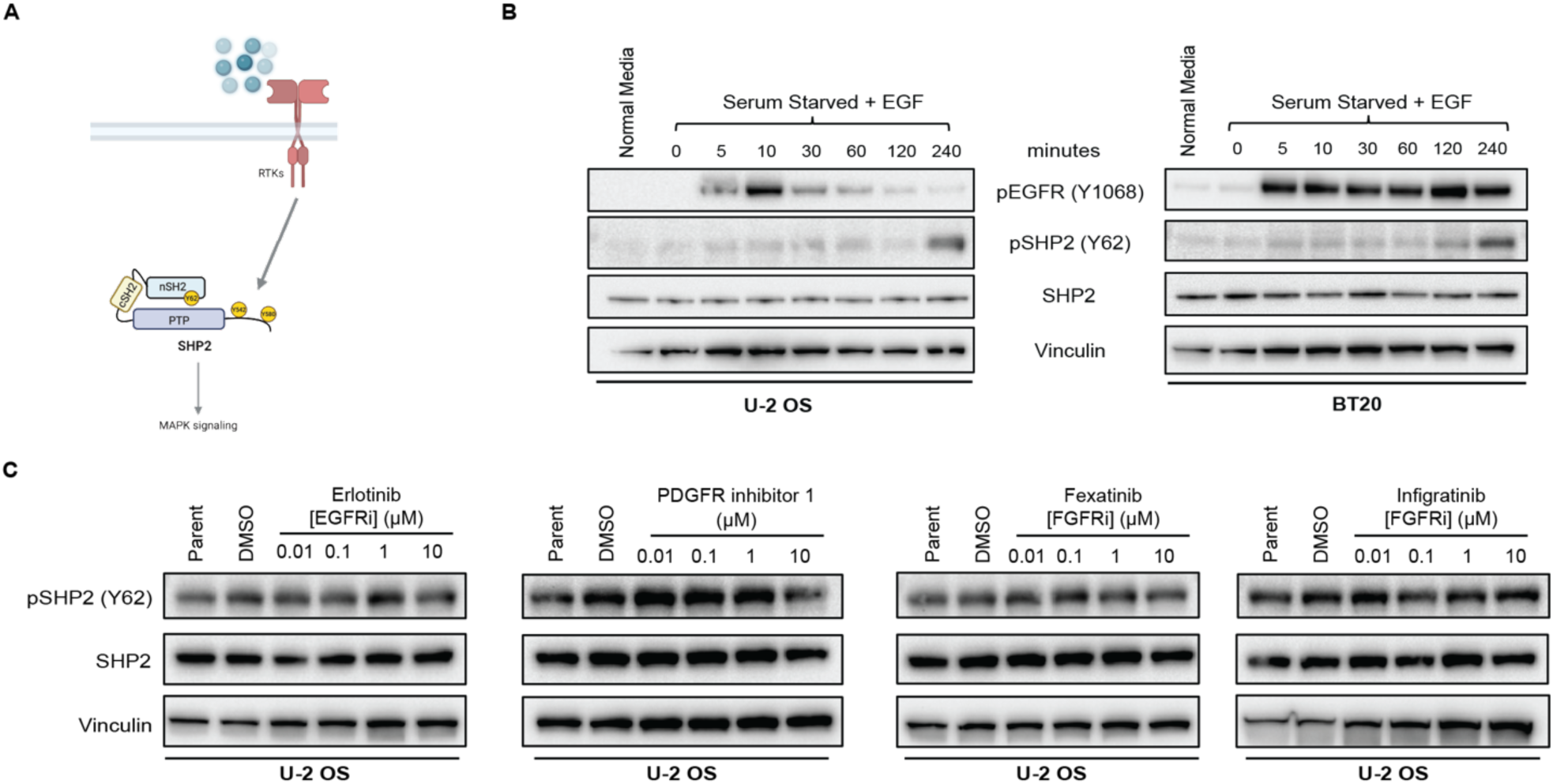
SHP2 pY62 is downstream of RTKs, but Y62 is not phosphorylated by RTKs. (**A)** RTKs signal to SHP2 through Y542 and Y580 phosphorylation. (**B**) Immunoblot analysis of U-2 OS and BT20 cells, serum starved (24 hours) and treated with EGF (100 ng/mL) for indicated timepoints. **(C)** Immunoblot analysis of U-2 OS cells treated with RTK inhibitors at indicated concentrations and DMSO. Erlotinib – EGFR inhibitor; Fexagratinib and Infigratinib – FGFR inhibitors; PDGFR inhibitor 1 – PDGFR inhibitor.

### SRC family kinases (SFKs) phosphorylate SHP2 Y62, Y542, and Y580

To determine the direct kinase for SHP2 Y62, we leveraged The Kinase Library, a recently developed global atlas of protein kinome in vitro substrate specificities^43,44^, based on the peptide primary sequence surrounding the phosphosite (**Supp. Data 3**). For SHP2 pY62, RTKs were predicted as the top candidate kinases (**Fig. 3A**). JAK3, a non-RTK, was also highly predicted. However, exposure of cells to the JAK3 inhibitor tofacitinib did not alter the level of SHP2 pY62 (**Supp. Fig. 3A**). Interestingly, non-RTKs, such as SRC family kinases (SFKs) SRC, YES1, and FYN, were in the bottom quartile with lower confidence scores. Since inhibiting RTKs did not alter the SHP2 Y62 phosphorylation, we wanted to test if SFKs can phosphorylate SHP2 Y62.

**Figure 3:**
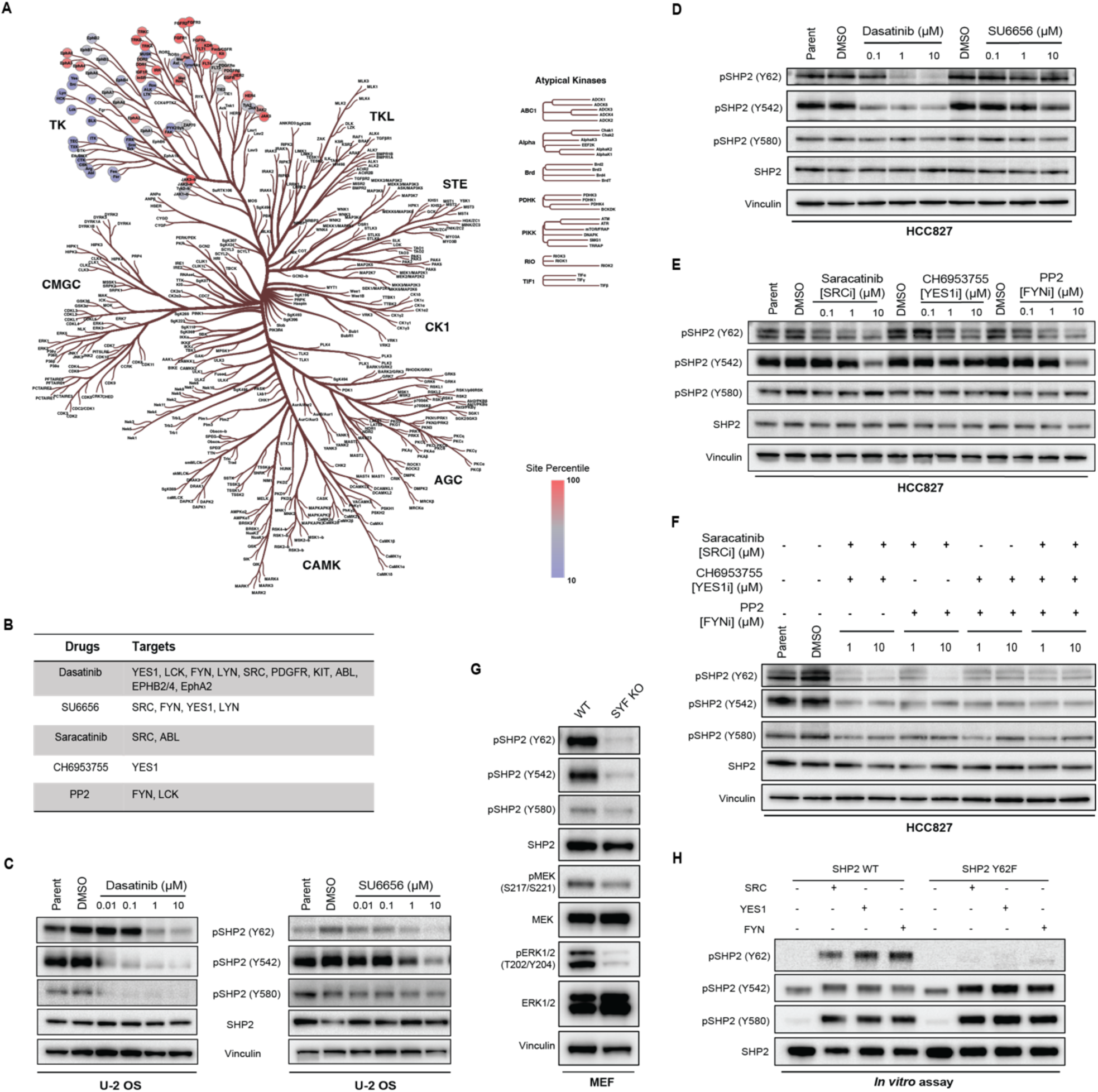
SRC, YES1, and FYN (SYF) kinases phosphorylate SHP2 Y62, Y542, and Y580. **(A)** Kinase Library prediction heatmap based on the peptide primary sequence of the SHP2 Y62 phosphosite plotted on human kinome evolutionary tree. (**B)** SFK multi-kinase and selective inhibitors with targets shown, tested here. (**C-D**) Immunoblot analysis of (**C**) U-2 OS cells and (**D**) HCC827 cells treated with dasatinib and SU6656 at indicated concentrations, and DMSO. (**E-F**) Immunoblot analysis of HCC827 cells treated with (**E**) saracatinib, CH6953755, and PP2, and (**F**) double and triple combinations at indicated concentrations, and DMSO. (**G**) Immunoblot analysis of SYF knock out and wildtype MEFs. (**H**) *In vitro* kinase assays with recombinant SRC, YES1, and FYN kinases and recombinant SHP2 WT and Y62F. The SHP2 pY542 antibody demonstrated nonspecific binding in the control conditions.

To test this hypothesis, we first used pharmacologic methods (**Fig. 3B**). Dasatinib is a multi-kinase inhibitor that targets not only SFKs but also ABL, PDGFR, and c-KIT^45^ (**Fig. 3B**). Treatment with dasatinib readily decreased SHP2 pY62 levels in U-2 OS cells (**Fig. 3C**) and HCC827 cells (EGFR-mutant NSCLC) (**Fig. 3D**). Dasatinib treatment also decreased SHP2 pY542 and pY580 levels in U-2 OS cells (**Fig. 3C**) but only pY542 in HCC827 cells (**Fig. 3D**). Since dasatinib has a wide range of targets including SFKs, ABL, and c-KIT, we next chose the selective SFK inhibitor SU6656^46^ which inhibits SRC, YES1, FYN, and LYN; asciminib^47^ which binds to the myristoyl binding pocket and inhibits ABL; and ripretinib^48^, which inhibits c-KIT, to narrow down candidate kinases (**Fig. 3B**). Treatment of U-2 OS cells with SU6656 decreased SHP2 pY62, pY542, and pY580 but there was no change in SHP2 phosphorylation with SU6656 in HCC827 cells (**Fig. 3D**). Treatment with asciminib or ripretinib did not change SHP2 phosphorylation levels (**Supp. Fig. 3B).**

The SFKs comprises 8 human kinases^49^, three of which–-SRC, YES1, and FYN–-are ubiquitously expressed in all tissues. In contrast, other SFK kinases (BLK, FGR, HCK, LCK, and LYN) are expressed in specific tissue types^50^. To our knowledge, there are no SFK inhibitors selective to individual SFKs. To better deconvolute the contributions of specific SFK kinases, we tested the SFK inhibitors saracatinib^51^ (SRC and ABL inhibitor), CH6953755^52^ (YES1 inhibitor), and PP2^53^ (FYN and LCK inhibitor) (**Fig. 3B**) which are more selective than dasatinib or SU6656 but less potent. In HCC827 cells, SHP2 Y62 phosphorylation was inhibited by saracatinib, CH6953755 and PP2, and Y542 phosphorylation was altered by PP2 and saracatinib at high concentrations, but not Y580 phosphorylation (**Fig. 3E**). In U-2 OS cells, SHP2 Y62 phosphorylation was inhibited by saracatinib only at high concentrations, while Y542 and Y580 phosphorylation were inhibited by saracatinib even at 1 µM (**Supp. Fig. 3C**), but not by CH6953755 (**Supp. Fig. 3D**) or PP2 (**Supp. Fig. 3E**).

We further investigated the cell-type specificity of SHP2 Y62 phosphorylation. Since 1) dasatinib decreased SHP2 pY62 levels in HCC827 and U-2 OS cells, 2) inhibitors of the non-SFK kinase targets of dasatinib did not decrease SHP2 phosphorylation, and 3) selective SFK inhibitors inhibited SHP2 pY62 levels in HCC827 but not in U-2 OS cells, we hypothesized that SHP2 Y62 phosphorylation is maintained by distinct SFK family members in a context-dependent manner, and that broad SFK inhibition is required to suppress SHP2 pY62 across diverse cell types.

To test this hypothesis, we tested double and triple combinations of the SFK selective inhibitors saracatinib, CH6953755, and PP2 in HCC827 and U-2 OS cells. Combining two drugs reduced SHP2 pY62, pY542, and pY580 levels more than single drugs, and combining all three drugs reduced levels more than combining two drugs (**Fig. 3F**, **Supp. Fig. 3F**).

We also used mouse embryonic fibroblasts (MEFs) from the SYF knock out mouse (SRC-/-, YES1 -/-, FYN -/-)^54^ as a genetic tool to test the hypothesis that SFKs phosphorylate SHP2 pY62. We found that SHP2 pY62, pY542, and pY580 levels were inhibited compared to WT MEFs (**Fig. 3G**). Thus, pharmacologic inhibition and genetic loss of the SFKs SRC, YES1, and FYN reduce SHP2 phosphorylation at Y62, Y542, and Y580.

Our initial growth factor experiments showed that SHP2 Y62 phosphorylation is downstream of RTKs (**Fig. 2B**). We used our pharmacologic and genetic models to situate SFKs in RTK-SHP2 signaling. EGF-dependent SHP2 phosphorylation at Y62, Y542, and Y580 was abrogated when U-2 OS cells were pretreated with dasatinib or SU6656 (**Supp. Fig. 3G**). EGF-, FGF1-, and PDGDF-dependent SHP2 phosphorylation at Y62 was abrogated in our SYF KO MEF models compared to WT MEFs (**Supp. Fig. 3H-J**). In contrast, SHP2 Y542 and Y580 phosphorylation were abrogated in SYF KO MEFs under FGF1 stimulation but not under EGF or PDGF stimulation (**Supp. Fig. 3H-J**).

Finally, to further validate that SFKs phosphorylate SHP2 Y62, we performed *in vitro* kinase assays using bacterially purified recombinant SHP2. SRC, YES1, and FYN kinases phosphorylated SHP2 WT at Y62 but not the Y62F phosphodead mutant (**Fig. 3H**). SRC, YES1, and FYN also phosphorylated Y542 and Y580 *in vitro* (**Fig. 3H**). Together these data show that SHP2 Y62 is a direct substrate of SFKs, and that SHP2 Y542 and Y580 can also be phosphorylated by SFKs.

### SHP2 Y62 phosphomimetic protein constitutively increases phosphatase activity

SHP2 Y62 phosphorylation has been proposed to disrupt its autoinhibited state through the introduction of negative charge at the N-SH2/PTP interface, thereby enhancing phosphatase activity, and this has been supported by biochemical and cellular experiments with the Y62 phosphomimetic mutants^30,55,56^. We corroborated these prior results by conducting Michaelis-Menten enzyme kinetics with the Y62D mutant, which showed a 2.4-fold increase in k_cat_/K_m_ when compared with wild-type SHP2 (**Fig. 4A**). Tyrosine to aspartate mutation is commonly used as a phosphomimetic to model the effects of phosphorylation, though it may not fully recapitulate the structural or binding consequences of a bona fide phosphotyrosine at this site. By contrast, the Y542 and Y580 phosphorylation sites are thought to mediate intramolecular interactions with the SHP2 SH2 domains, or mediate intermolecular interactions with other SH2-containing proteins. SH2-mediated interactions with tyrosine-phosphorylation proteins cannot be readily mimicked using D/E substitutions, and indeed, the Y542D and Y580D mutations did not significantly impact SHP2 activity (**Fig. 4A**). Furthermore, the combination of Y542D or Y580D with Y62D did not appreciably impact the gain-of-function effect of Y62D (**Fig. 4A**).

**Figure 4:**
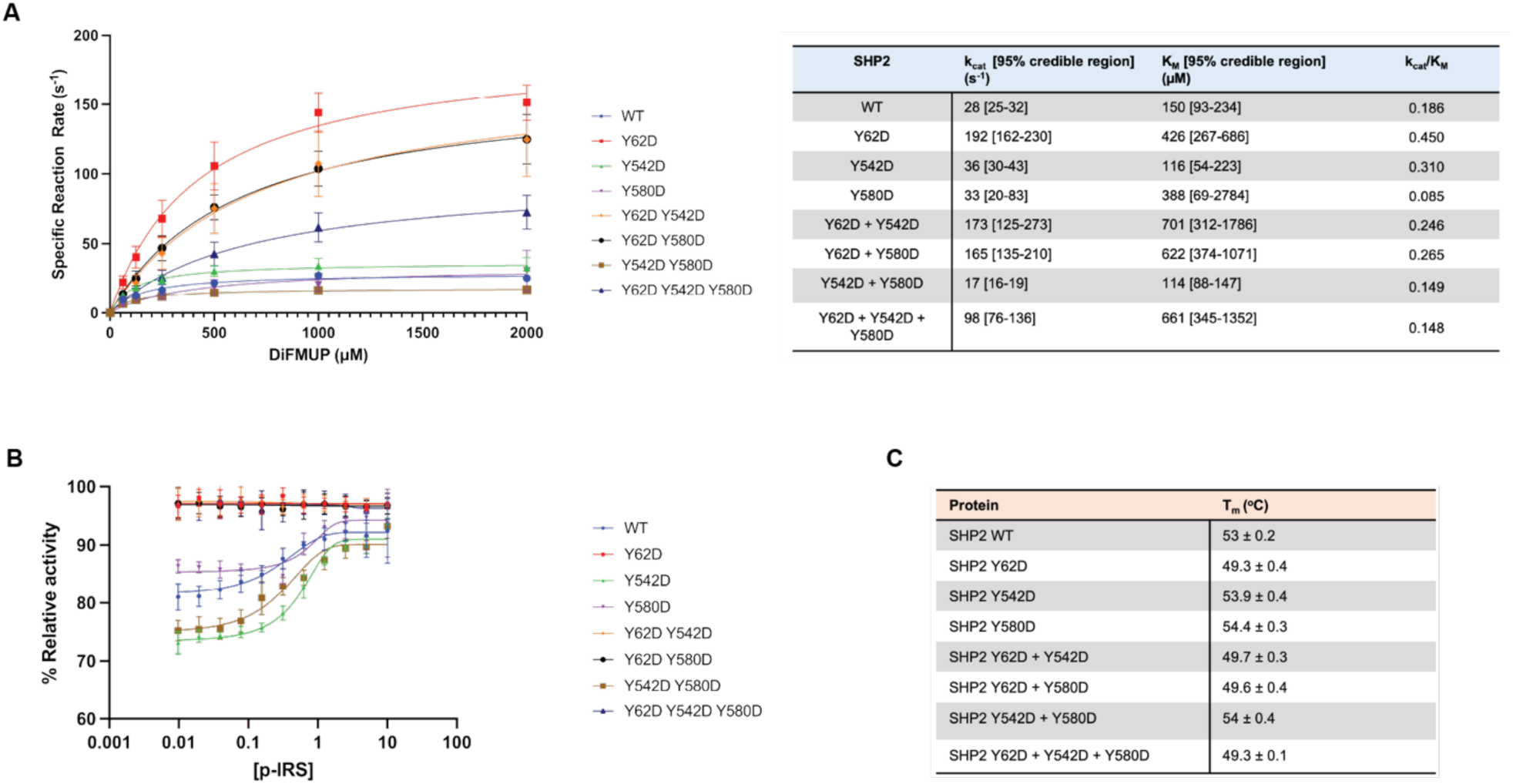
SHP2 Y62 phosphomimetic constitutively increases phosphatase activity. (**A)** Phosphatase enzyme kinetic reaction curves of wild-type and mutant SHP2 proteins across varying concentrations of DiFMUP. Catalytic constant (k_cat_) and Michaelis constant (K_m_) values are denoted in the table. (**B)** Phosphatase activity of SHP2 proteins across varying concentrations of an IRS-1-derived monophosphorylated peptide. (**C)** Melting temperatures (T_m_) of wild-type and mutant SHP2 proteins by differential scanning fluorimetry (DSF).

When SHP2 auto-inhibition is disrupted by mutations, it becomes less responsive to activation by phosphopeptides^57^. Indeed, unlike wild-type SHP2, Y62D-containing SHP2 variants could not be activated by binding to an IRS-1-derived phosphopeptide (**Fig. 4B**). To further examine whether these mutations disrupt auto-inhibition, we used differential scanning fluorimetry (DSF) (**Supp. Fig. 4**). Previous studies have shown that SHP2 variants with disrupted autoinhibition tend to have lower melting temperatures^58^. Indeed, only SHP2 variants containing the Y62D mutation, but not Y542D or Y580D, had decreased melting temperatures (**Fig. 4C**). Collectively, these experiments show that the Y62D mutation disrupts SHP2 auto-inhibition and corroborates the assertion that Y62 phosphorylation may disrupt auto-inhibition through electrostatic effects.

### SHP2 Y62D samples mixed open and closed conformations through increased exchange in its buried nSH2 domain

Since the SHP2 Y62D phosphomimetic protein exhibited increased enzymatic activity (**Fig. 4A**) and decreased thermal instability (**Fig. 4C**), and since SHP2 oncogenic mutation propensity for an open conformation correlates with increased catalysis and increased dynamic and solvent exposure^59,60^, we hypothesized that SHP2 Y62 phosphorylation increases catalysis by enforcing an open, dynamic conformation. We investigated this hypothesis using computational, structural, and biophysical methods. We used AlphaFold2 to analyze conformational changes in SHP2 Y62D and compared the predicted structure to crystal structures of SHP2 wild-type (WT), which adopts a closed conformation (PDB: 2SHP)^8^, and SHP2 E76K, the only SHP2 mutant with a crystal structure resolved in an open conformation (PDB: 6CRF)^57^. While AlphaFold2 is not specifically powered for optimal structural prediction of missense mutations, nonetheless, SHP2 Y62D was predicted to have a 22 Å root mean square deviation (RMSD) when aligned with the WT crystal structure, but a 1.1 Å RMSD when aligned with the SHP2 E76K structure oncogenic mutant crystal structure, suggesting an open confirmation (**Fig. 5A**). We crystallized the SHP2 Y62D mutant and solved its structure at 2.9 Å resolution (**Fig. 5B, Supp. Table 1**). The Y62D crystal structure exhibited a closed conformation, with a 0.5 Å RMSD when aligned with the SHP2 WT crystal structure (**Fig. 5B**).

**Figure 5:**
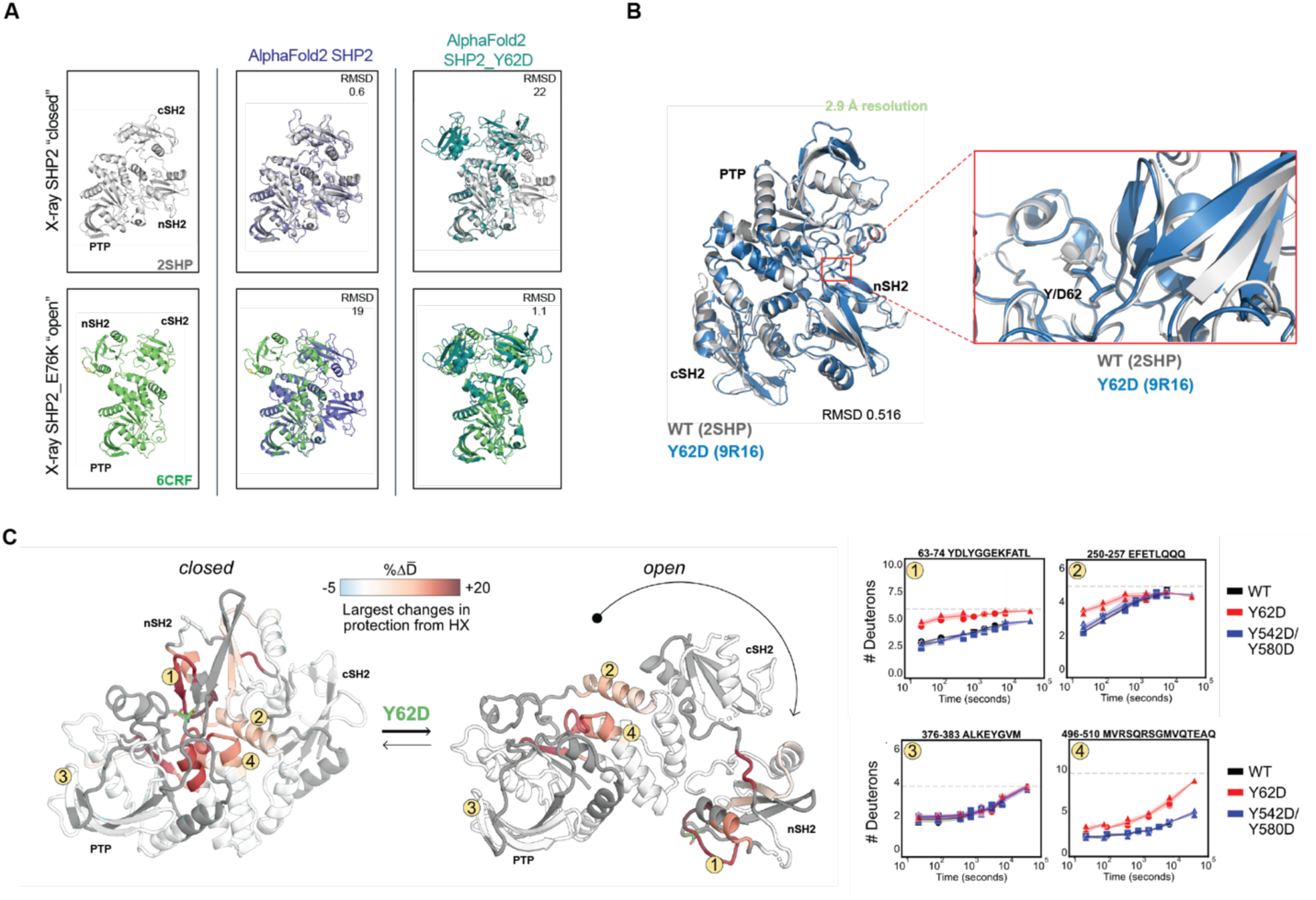
SHP2 Y62D samples both the closed and open state conformations through increased exchange in its buried nSH2 domain. **(A)** Superimpositions of AlphaFold2-predicted structures of SHP2 Y62D mutant (sky blue) and WT (purple) onto crystal structures of SHP2 E76K (green) (PDB: 6CRF) and WT (gray) (PDB: 2SHP) with root mean square deviation (RMSD) values indicated. **(B)** Crystal structure SHP2 Y62D mutant (blue) (PDB: 9R16) superimposed onto SHP2 WT crystal structure (gray) (PDB: 2SHP). **(C)** Peptides that show hydrogen-deuterium exchange mass spectrometry (HDX-MS) differences plotted onto SHP2 WT crystal structure (closed conformation) (PDB: 2SHP) and SHP2 E76K crystal structure (open conformation) (PDB: 6CRF), with representative peptides indicated. HDX rates between Y62D and WT for the 63-74 and 496-510 peptides are different, while for the 376-383 peptide are similar. HDX rates between Y542D/Y580D and WT are similar for all peptides.

Given the discordance between our biochemical and computational data and compared to our structural data, we hypothesized that SHP2 Y62 phosphorylation samples both the closed and open state conformations. To investigate the conformational state of the SHP2 Y62D mutant, we performed hydrogen-deuterium exchange mass spectrometry (HDX-MS) with recently developed peptide disambiguation and analysis tools^61^ comparing SHP2 Y62D to wild-type (WT) SHP2 and the SHP2 phosphomimetic mutant Y542D/Y580D. Compared to WT, SHP2 Y62D exhibited increased deuterium uptake across peptides spanning not only the nSH2 domain (where Y62 is located), but also in the PTP domain mediated by amino acids that bind to and maintain nSH2 domain autoinhibitory interactions (**Fig. 5C, Supp. Fig. 5**). Notably this PTP domain peptide region (496-510) also includes S502, G503, and Q506 which bind to the nSH2 domain, are known cancer-associated mutations^60^, and activate SHP2 in deep mutational scanning studies^62^. This pattern is consistent with increased destabilization of the closed, autoinhibited conformation and adoption of an open conformation relative to WT SHP2. In contrast, the Y542D/Y580D mutant showed minimal differences in hydrogen exchange relative to WT, indicating that these phosphomimetic substitutions do not strongly alter the conformational ensemble (**Supp. Fig. 5**). Together, these data demonstrate that the Y62D mutation promotes an open, active-like conformation of SHP2, distinct from WT and Y542D/Y580D.

### SHP2 Y62 phosphomimetic mutation activates MAPK signaling and causes resistance to SHP2 inhibitors

Since SHP2 pY62 is enriched in RTK-driven tumors, we tested the role of SHP2 Y62 phosphorylation in cancer cells. We knocked out endogenous *PTPN11*/SHP2 using CRISPR-Cas9 in BT20 and U-2 OS cells, which are frequently used to model SHP2 alterations^30,57^ (**Supp. Fig. 2A**), and complemented this with lentiviral overexpression of SHP2 WT, aspartate and phenylalanine mutants at Y62 and Y542/Y580. Complementation with empty vector control was not tolerated by cells, consistent with *PTPN11*/SHP2 loss being synthetic lethal in RTK-driven cancer cells^20^. SHP2 Y62D, but not Y62F, sustained maximal MAPK signaling as measured by phosphorylated MEK (pMEK) and phosphorylated ERK (pERK) in SHP2 KO BT20 cells, suggesting that SHP2 pY62 is necessary for maximal MAPK signaling (**Fig. 6A**). To delineate how upstream SFKs modulate MAPK signaling, we used our SYF KO MEF models. SYF KO showed decreased pMEK and pERK as compared to WT MEFs (**Fig. 3G**).

**Figure 6:**
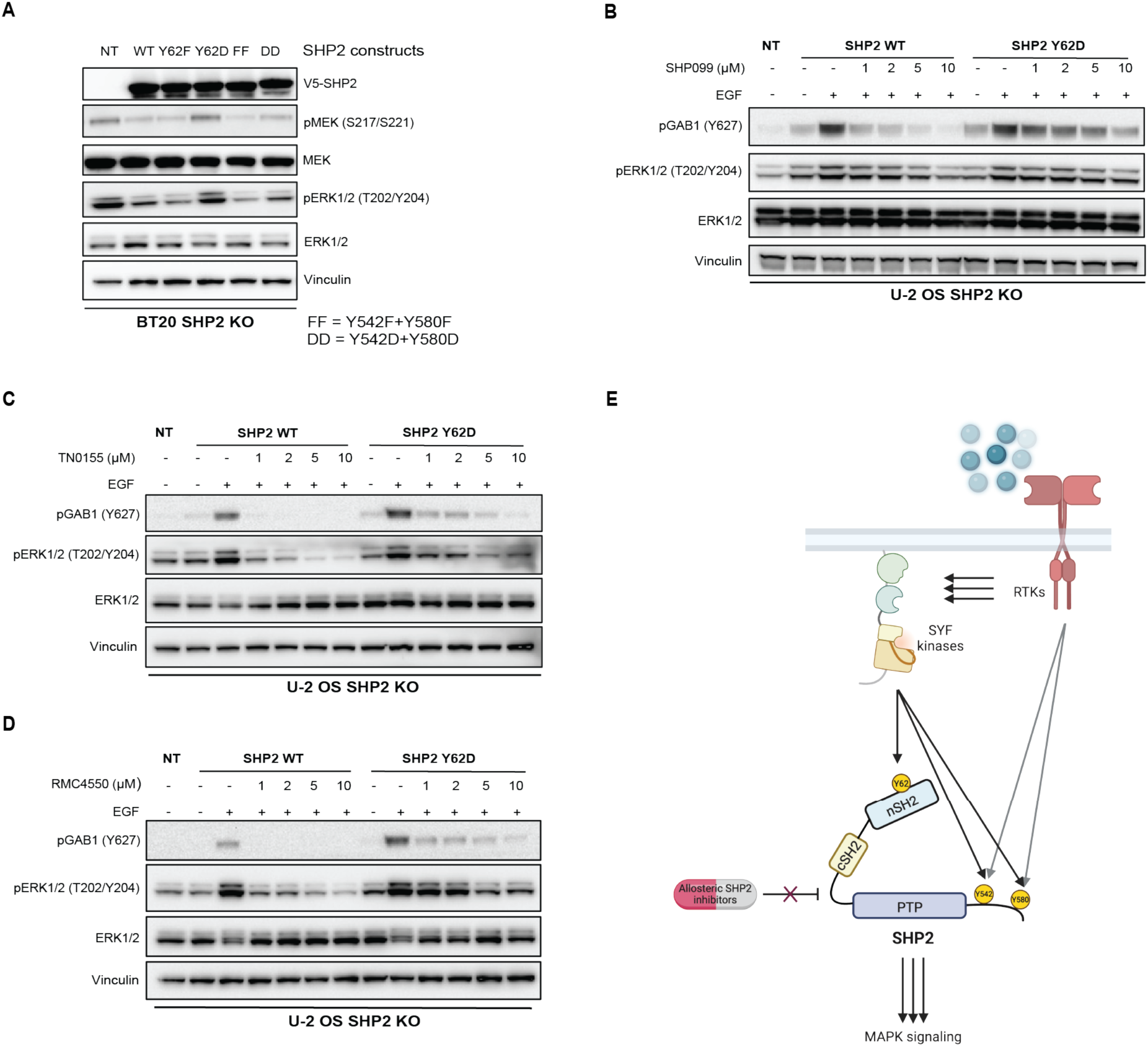
SHP2 Y62 phosphomimetic mutation causes resistance to SHP2 inhibitors. (**A)** Immunoblot analysis of BT20 SHP2 knock out cells expressing SHP2 mutants. FF, Y542F/Y580F; DD, Y542D/Y580D. (**B**-**D**) Immunoblot analysis of U-2 OS SHP2 knock out cells expressing SHP2 WT and SHP2 Y62D proteins, serum starved then treated with (**B**) SHP099, (**C**) TN0155, and (**D**) RMC4550 followed by EGF (100 ng/mL) treatment for 10 minutes. (**E**) Model for SHP2 Y62 phosphorylation by SYF kinases, conformational changes, and drug resistance.

Given the abundance of SHP2 pY62 across cancer cells and tumors, and the minimal response rates conferred by SHP2i in the clinic, we hypothesized that SHP2 pY62 confers resistance to SHP2i. We tested varied allosteric SHP2 inhibitors including SHP099, RMC4550, and TNO155, which are based on similar scaffolds that stabilize the SHP2 autoinhibited conformation^20,63,64^. SHP2 Y62 phosphorylation as modeled by SHP2 Y62D resulted in resistance to SHP099, RMC4550, and TNO155, as measured by sustained pERK signaling, as well increased GAB1 phosphorylation which is a direct readout of SHP2 scaffolding function^65^ (**Fig. 6B-D**).

## Discussion

We elucidated a SHP2 pY62 hotspot phosphorylation catalyzed by SFK, which functionally phenocopies the known rare activating mutations of *PTPN11*/SHP2, yielding a constitutively active phosphatase, enrolling an active, open state potentiating oncogenesis and SHP2 inhibitor resistance (**Fig. 6E**).

SHP2 was first discovered as a tyrosine phosphatase oncoprotein phosphorylated in response to growth factors^1–5^. Significant investigations into the consequences of RTK-mediated phosphorylation at SHP2 Y542 and Y580 over the decades have shown that they enable SHP2 to recruit signaling complexes for downstream signaling^15,25–27^. The precise contribution of these phosphosites, located in the unstructured C-terminal tail, in SHP2 catalytic activation has been debated^28^. These sites mediate response to some but not all growth factors, raising the idea of alternative regulatory mechanisms for SHP2.

In its initial discovery, SHP2 was also ubiquitously tyrosine-phosphorylated in cells transformed by the v-*src* retroviral oncogene^1^. However, whether human SRC (or other SFK family kinases) directly phosphorylate and regulate SHP2 has remained unknown. Our findings implicate SHP2 Y62 as the site of SHP2 regulation by SRC and other SFKs. Moreover, our findings suggest a unifying mechanism for kinase phosphorylation and SHP2 activation, where multiple phosphorylation inputs drive SHP2 into an active state: by C-terminal phosphorylation increasing scaffolding phosphotyrosine-dependent recruitment and by nSH2 domain modification as we describe. Notably, there is significant cell-type heterogeneity as to which specific RTKs and SFKs dominate in the RTK-SFK-SHP2-MAPK signaling axis we have delineated.

SHP2 pY62 has been documented previously in isolated mass spectrometry datasets of drug-treated lung cancer and leukemia cell lines^30,55^. Supporting the importance of this site, a rare Y62D germline mutation in Noonan syndrome patients has also been described^56^. That SHP2 pY62 is highly prevalent in solid tumors, despite the rarity of *PTPN11* mutations, indicate that SHP2 activation in solid tumors occurs predominantly by post-translational modification or through phosphopeptide binding rather than genomic alterations, which are more commonly seen in hematologic malignancies.

These findings have important therapeutic implications. Allosteric SHP2 inhibitors currently in clinical development have demonstrated minimal efficacy, with objective response rates of ∼0% in phase 1 clinical trials, and no resistance mechanisms have been identified to date. Our data implicate intrinsic SHP2 Y62 phosphorylation as a primary resistance mechanism by stabilizing SHP2 in an open, active conformation. These results provide a mechanistic rationale for the limited clinical activity observed with allosteric SHP2 inhibitors and indicate that elevated SHP2 pY62 could serve as a biomarker of intrinsic resistance. Furthermore, they identify Y62-phosphorylated SHP2 as a structurally and dynamically distinct target from WT SHP2, necessitating alternative strategies for mutant selective or phospho-selective inhibition.

## Methods and Materials

### Data reproducibility statement

To ensure the rigor and reproducibility of our findings, all experimental procedures were conducted following standardized protocols with appropriate controls. Detailed methods, including reagent sources, catalog numbers, and RRIDs, are provided to enable exact replication. Biological and technical replicates were performed to confirm consistency, and statistical analyses adhered to established guidelines.

### In silico analysis of pY62 abundance

Phosphorylation_site_dataset.gz was obtained from PhosphoSitePlus^31^. MS_LIT (the number of mass spec studies supporting the site) was added to MS_CST (the number of mass spec studies performed by Cell Signaling Technology supporting the site) to obtain the number of observations. LT_LIT (the number of publications supporting the site) was divided by the total number of observations ((LT_LIT+1)/(MS_CST+MS_LIT) to obtain the ratio of literature support and abundance of phosphopeptide observation.

The phosphosite abundance in KRAS mutations vs WT tumors and EGFR mutations vs WT tumors in lung adenocarcinoma was obtained from Gillette et al (2020) – Supplementary table S4L and S4M^38^. The phosphosite abundance in FGFR altered vs non-altered tumors in intrahepatic cholangiocarcinoma was obtained from Dong et al (2022)^39^. Datasets were plotted with GraphPad (Prism, RRID:SCR_002798).

### Cell lines used in this study

HEK293T LentiX (ATCC - CRL-3216, RRID:CVCL_0063), U-2 OS (ATCC - HTB-96™, RRID:CVCL_0042), BT20 (ATCC - HTB-19™, RRID:CVCL_0178), HCC827 (ATCC - CRL 2868, RRID:CVCL_2063), SYF KO (ATCC - CRL-2459, RRID:CVCL_6461), and SV40 immortalized Mouse embryonic fibroblast – MEFs (Gift from Assistant Prof. Iok In Christine Chio, Columbia University) were used in this study and frequently checked for mycoplasma contamination. U-2 OS, SYF KO MEFs, and WT MEFs were maintained in DMEM media (Gibco, Cat# 11965-092) with 10% FBS (Sigma, Cat# F0926) and 1% Pen Strep (Gibco, Cat# 15070-063). BT20 was maintained in DMEM/F12 media (Gibco, Cat# 11330-032) with 10% FBS and 1% Penicillin/Streptomycin. HCC827 was maintained in RPMI media 1640 (Gibco, Cat# 11875-093) with 10% FBS and 1% Pen Strep. TrypLE™ Express (Gibco, Cat# 12604-021) was used to detach the cells from the plates.

### RTK stimulation

The stock solution (100mg/mL) of Human EGF (Sigma-Aldrich, Cat# E9644), FGF1 (SignalChem, Cat# F819-40N-50), and PDGF-ββ (Sigma-Aldrich, Cat# SRP3138) were made with ddH_2_O and stored at -20°C. The cells were serum starved for 24 hours prior to the RTK stimulation. The ligands were diluted in appropriate media and spiked in the cells. The stimulation was stopped at different time points by washing with ice cold PBS wash.

### Drug treatment

All drugs were diluted with DMSO to acquire 10mM stock concentration and were diluted with appropriate media prior to the drug treatments. Erlotinib (Cat# S7786), PDGFR inhibitor 1 (Cat# S8721), Tofacitinib (Cat# S2789), Dasatinib (Cat# S1021), Saracatinib (Cat# S1006), SHP099 (Cat# S6388), TN0155 (Cat# S8987), and RMC4550 (Cat# S8718) were purchased from Selleckchem, Houston, TX, USA. Fexagratinib (Cat# HY-13330), Infigratinib (Cat# HY-13311), Asciminib (Cat# HY-104010), Ripretinib (Cat# HY-112306), SU6656 (Cat# HY-B0789), CH6953755 (Cat# HY-135299), and PP2 (Cat# HY-13805) were purchased from MedChemExpress, NJ, USA. For drug treatment assays (RTK inhibitors and SFK inhibitors), the cells were treated with final concentrations of the desired drugs in the normal media. For EGF stimulation in the presence of dasatinib and SU6656, Suppl. Fig 3G, the cells were serum starved for 24 hours, followed by an hour drug treatment and RTK stimulation at different timepoints. For SHP2 inhibitors treatment with EGF stimulation, the cells were serum starved for 24 hours and treated with allosteric SHP2 inhibitors for an hour and followed by 10 minutes EGF stimulation (final concentration 100ng/mL). The reactions were stopped by washing with ice cold PBS wash.

### Western blotting analysis

Cells were lysed in Pierce™ RIPA lysis buffer (Thermo Scientific, Cat# PI87788) containing protease (Selleckchem, Cat# B14001) and phosphatase inhibitors (Selleckchem, Cat# B15001). BCA assay (Thermo Scientific, Cat# 23225) was performed to determine the protein concentration. Samples were run on PAGE gels (ThermoFisher) and transferred to Trans-Blot transfer membranes (Bio-Rad, Cat# 1620115) using the Trans-Blot® Turbo™ Transfer System. The membranes were blocked with 5% milk in 1x TBST (10x TBS - Bio-Rad, Cat# 1706435; Tween® 20- Fisher, Cat# 240186) and probed with the following primary antibodies (pEGFR- Y1068 (1:1000, Cat# 3777S, RRID:AB_2096270), pFGFR-Y653/654 (1:1000, Cat# 3471S, RRID:AB_331072), pPDGFR-Y751 (1:1000, Cat# 3161S, RRID:AB_331053), pGab1-Y627 (1:1000, Cat# 3233S), pSHP2-Y62 (1:500, This study), pSHP2-Y542 (1:1000, Cat# 3751S, RRID:AB_330825), pSHP2-Y580 (1:1000, Cat# 3703S), SHP2 (1:1000, Cat# 3397S, RRID:AB_2174959), pMEK-S217/S221 (1:1000, Cat# 9154S), MEK (1:1000, Cat# 8727S), pERK-T202/Y204 (1:1000, Cat# 4370S), ERK (1:1000, Cat# 4696S), Vinculin (1:1000, Cat# 13901S, RRID:AB_2728768), and V5 (1:1000, Cat# 13202S, RRID: AB_2687461) from Cell Singaling Technologies) and secondary antibodies (anti-Mouse IgG (1:4000, Cat# 7076S, RRID: AB_330924), and anti-Rabbit IgG (1:4000, Cat# 7074S, RRID: AB_2099233) from Cell Singaling Technologies] before developing with SuperSignal West Atto Ultra-Sensitivity Chemiluminescent Substrate (ThermoFisher: A38556). Blots were detected using the ChemiDoc MP Imaging System (Bio-Rad). All experiments were performed in triplicate.

### Protein expression and purification

cDNA of SHP2 (UNIPROT ID Q06124) was ordered as a Genestring and cloned into a pGEX-5X- 1 vector (RRID: Addgene_29352) with a N-terminal GST tag and thrombin cleavage site for in vitro experiments. For all other in vitro, the cDNA was cloned into a pHAT vector (RRID: Addgene_112582) with a non-removable N-terminal his-tag. For lentiviral overexpression, the cDNA was cloned into pLX304 (RRID:Addgene_25890) with a C-terminal v5 tag. pGEX-5X-1_SHP2C495S, pGEX-5X-1_SHP2Y62F, pHAT_SHP2Y62D, pHAT_SHP2Y542D, pHAT_SHP2Y580D, pHAT_SHP2Y62D/Y542D, pHAT_SHP2Y62D/Y580D, pHAT_SHP2Y542D/Y580D, pHAT_SHP2Y62D/Y542D/Y580D, pLX304_SHP2Y62F, pLX304_SHP2Y62D, pLX304_SHP2Y542F/Y580F, pLX304_SHP2Y542D/Y580D were prepared through site directed mutagenesis, designed with NEBaseChanger.

pGEX-5X-1_SHP2 or pHAT_SHP2 constructs were transformed into electrocompetent *E. coli* BL21(DE3) (NEB, Cat# C2527H). All constructs were expressed in 0.5-1L LB with 100 µg/mL carbenicillin/ampicillin, induced at OD600 0.4-0.6 with 0.1 mM IPTG and grown overnight at 20 °C. Cell pellets were washed with binding buffer (20 mM Tris, 500 mM NaCl, 5 mM Imidazole, pH 8.0) and spun down.

For pHAT_SHP2 constructs, the washed cells were resuspended with 10-15 mL lysis buffer (binding buffer, 1x Bugbuster (Millipore, Cat# 70921), 1 tablet protease inhibitor/50 mL (Pierce, Cat# A32963), 1 mg/mL Lysozyme from chicken egg white (Sigma, Cat# L8676), 0.2 uL/mL nuclease (Pierce, Cat# 88701) and lysed for 30 minutes at room temperature. The supernatant was separated (15000g, 30 min) and added to 1.5 mL CV of nickel resin (His-select nickel affinity gel, Merck, P6611 on a gravity-based column. Nickel resin was washed with 5 CV binding buffer with 2 mM TCEP before being eluted with 2.5 mL elution buffer (20 mM Tris, 500 mM NaCl, 500 mM Imidazole, 2 mM TCEP pH 8.0). The eluted protein was dialyzed with PD10 columns (Cytiva, Cat# 17085101) to SHP2 buffer (25 mM Tris-HCl, 150 mM NaCl, 2 mM DTT, pH 8.0). The dialyzed protein was size excluded on a Superdex SD75 column (Cytiva, Cat# 10/300 GL) and concentrated through Amicon centrifugal filters (Millipore, Cat# UFC803024, 4 mL, 30K). All proteins were aliquoted for single-use, flash-frozen, and stored at -80 °C.

For pGEX-5X-1_SHP2 constructs, the washed cells were resuspended with 10-15 mL lysis buffer (PBS (Gibco, Cat# 20012-027), 1x Bugbuster, 1 tablet protease inhibitor/50 mL (Pierce, Cat# A32963), 1 mg/mL Lysozyme from chicken egg white (Sigma, Cat# L8676), 0.2 uL/mL nuclease (Pierce, Cat# 88701) and lysed for 30 minutes at room temperature. The supernatant was separated (15000g, 30 min) and combined with 2 mL CV glutathione resin (Pierce Glutathione Agarose, Cat# 16101) washed with equilibration buffer (50 mM Tris, 150 mM NaCl, pH 8.0) rotating overnight at 4 °C. The resin-supernatant mixture was added to a gravity column and the flow through was discarded. The resin was washed by resuspending the resin with 5 mL equilibration buffer. Thrombin (Sigma-Aldrich, Cat# T4648-1KU) resuspended in 1 mL cold PBS (1U/μL), was diluted 10x in equilibration buffer, and 100 units of thrombin (1 mL) were added to the column, cleaving overnight at 4 °C. One mL of protease inhibitor solution (Pierce, Cat# A32963, one tablet/50 mL) was added to the column, collecting the eluted fraction. The eluted fraction was dialyzed to RTK buffer (20 mM Tris, 300 mM NaCl, 1 mM TCEP, 5 mM MgCl2, 20% glycerol, pH 8.0) after which ∼5 v/v% of concentrated protease inhibitor was added (1 tablet protease inhibitor/mL, Pierce, Cat# A32963). All proteins were aliquoted for single-use, flash-frozen and stored at -80 °C.

### *In vitro* kinase assay

SRC (Cat# S19-18G), YES1 (Cat# Y01-10G), and FYN (Cat# F15-10G) kinase proteins were purchased from SinoBiological. Reaction mixture was prepared with the 5x Kinase buffer (Cell Signaling Technology, Cat# 9802S), ATP (Cell Signaling Technology, Cat# 9804S, final concentration 1 mM), SHP2 recombinant protein (1 µg), the kinase of interest (0.2 µg) and ddH2O to make the final volume of 50 µL. The reaction mixture was incubated at 30C in Thermocyler for 1 hour and stopped by adding 4x sample buffer (Life Technologies, Cat# NP0008) in the reaction mixture and incubation at 95C for 5 mins.

### Michaelis-Menten kinetics

Phosphatase activity was measured using the EnzChek® Phosphatase Assay kit for SHP2 activity (Invitrogen, Cat# E12020) according to the manufacturer’s protocol. Briefly, the reactions were initiated by the addition of 0-2000 µM 6,8-Difluoro-4-Methylumbelliferyl Phosphate (DiFMUP) to the reaction mixture containing purified SHP2 variants in 1x reaction buffer (100 mM sodium acetate (pH 5.0). The fluorescence was measured using excitation at 360 nm and emission was recorded at 455 nm with a Tecan Infinite M200 Pro Plate Reader. Initial reaction rates were determined and divided by the final enzyme concentration to determine specific reaction rates. The specific reaction rates for each SHP2 variant were plotted against substrate concentration and the curves were fitted to the Michealis-Menten equation using GraphPad (Prism, RRID:SCR_002798) to establish *k*_cat_ and *K*m values.

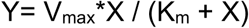

Where X is substrate concentration, Y is enzyme velocity, V_max_ is maximum reaction rate and K_m_ is Michaelis constant

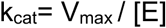

Where k_cat_ is turnover number and [E] is total enzyme concentration

### p-IRS stimulation

p-IRS (GLN{pTyr}IDLDLVKD, GenScript, Lot# U644VGB170-1/PE4713) was resuspended in DMSO to a concentration of 3 mg/mL. SHP2 variants (0.15 μM final, in SHP2 buffer) were combined with a linear titration of p-IRS in HEPES buffer (10 μM-0.01 μM final) and incubated for 30 minutes at room temperature. DiFMUP (100 μM final) was added to start the reaction, and fluorescence was measured at t = 30 min by the Tecan Infinite M200 Pro Plate Reader (RRID:SCR_019033).

### T_m_ measurements

SHP2 variants were diluted to 2 mg/mL in SHP2 buffer (25 mM Tris-HCl, 150 mM NaCl, 2 mM DTT, pH 8.0). SYPRO orange (5000x, Thermo Fisher) was diluted to a 25x solution in SHP2 buffer. In a 20 uL reaction, 5 uL SHP2 (0.5 mg/mL final), 4 uL SYPRO orange (5x final) was combined in SHP2 buffer in triplicate. The sample was measured on an Applied Biosystems 7500 Fast Real-Time PCR (RRID:SCR_018051), using ROX as a reporter with a 0.1 °C/sec slope from 25 to 99°C. Melting temperature (Tₘ) was recorded at the inflection point of fluorescence vs. temperature curve.

### Crystallography

#### • Protein Production

The SHP2^Y62D^ construct used for crystallization comprises residues 1-528 of SHP2, the mutation Y62D and a non-cleavable C-terminal hexahistidine tag. The protein was produced in *E. coli* and purified by IMAC, ion-exchange chromatography and size-exclusion chromatography as described previously^66^.

#### • Crystallization

SHP2^Y62D^ was crystallized by hanging-drop vapor-diffusion at 20°C. 1.0 µL of SHP2^Y62D^ (13.7 mg/mL in 50 mM NaCl, 20 mM TRIS-HCl pH 7.8, 1 mM EDTA and 2 mM DTT) was mixed with 1.0 µL of crystallization solution (0.2 M BICINE pH 8.5 and 17% w/v PEG 4000) and equilibrated against a reservoir containing crystallization solution. The crystals were mounted after three days. Crystals were cryo-protected in crystallization solution supplemented with 20% v/v Glycerol (final concentration) and cooled in liquid nitrogen. Data were collected at beamline BL18U1 of the Shanghai Synchrotron Radiation Facility (Shanghai, China).

#### • Structure determination

Diffraction data were integrated, analyzed and scaled with *XDS*^67^*, POINTLESS, AIMLESS* and *STARANISO*^68^ in *AUTOPROC*^69^. The structures were determined by molecular replacement with *PHASER* using a previously determined model of SHP2 (without any ligands) as a starting model. The model was improved through manual rebuilding of the model in *COOT*^70^ and restrained refinement with *BUSTER*^71^. Atomic displacement factors were modelled with a single TLS group per chain and a single isotropic B-factor per atom. The backbone geometry was analyzed with *MOLPROBITY*^72^.

### HDX/MS Protocol

H_2_O buffer (25mM Tris-HCl, 150mM NaCl, 2mM DTT) was prepared in deionized water and lyophilized using Labconco FreeZone 4.5 lyophilizer in 5ml aliquots for at least 24 hours, then resuspended in D_2_O to produce deuterated buffer. HX liquid handling was performed using the LEAP system (Trajan) controlled by Chronos 5.8.3 software (Trajan). Protein stock integrity was verified by MALDI-MS (Bruker Autoflex Speed TOF/TOF). The protein stock solution was manually pipetted into reaction vials (to avoid syringe clogging), diluted tenfold in deuterated buffer (7.5µl protein stock in 75µl buffer), and exchanged at 15°C for the specified time, then diluted twofold (75µl exchange reaction) in a quench solution (75µl, 3% acetonitrile, 1% formic acid in H_2_O, all MS-grade, Fisher Scientific) at 2°C. 140 μL of the quenched reaction was injected into the protease column, which was chilled to 7 °C. Solvent A (3% acetonitrile, 0.15% formic acid in H_2_O) was pumped (UltiMate 3000, ThermoFisher Scientific) at 150 μL/min through the protease column. Peptides eluted from the protease column were trapped on a 1x10 mm C18 column (Hypersil GOLD, 3 μM poresize, ThermoFisher Scientific). For Rep1 and Rep2, we used an immobilized protease type XIII/pepsin column (FP, 1:1 w/w, 2.1 × 30 mm, NovaBioAssays), for Rep3 we used an alanyl aminopeptidase/pepsin column (AP/pepsin 2.1 x 20 mm, Affipro).

Fully deuterated sample buffer was prepared by adding 3M GuHCl to deuterated buffer. Full-D control samples were prepared by manually diluting protein stock 1:10 in Full-D buffer and exchanging at 37° for at least 24 hours. Exchanged Full-D reaction was pipetted into reaction vials (82.5µl per vial) and subsequently quenched and injected exactly as for other samples.

An auxiliary pump (MX-Class, Teledyne) flowed additional Solvent A over the trap C18 at 150 μL/min. After digesting and desalting (3min for rep1, 200s for rep2), the system valve was switched so that the binary pump (Infinity II, Agilent) was in-line with the trap C18, flowing in the opposite direction. The binary pump used a gradient of Solvent A and Solvent B (96.85% acetonitrile, 0.15% formic acid) to elute peptides from a 50 x 1 mm C18 analytical column (Hypersil GOLD, 1.9 μM pore size, ThermoFisher Scientific). The elution method consisted of isocratic flow at 10% B for 1 min, a linear gradient to 40% B from 1 to 15 min, a linear gradient to 95% B from 15 to 19 min, isocratic flow at 100% B from 19 to 24 min, a linear gradient to 10% B from 24 to 30 min, and then isocratic flow at 10% B from 30 to 45 min, at a flow rate of 0.04 mL/min. The protease column was washed between sample injections with injections of quench solution.

Eluted peptides entered the mass spectrometer (Bruker Maxis II ETD) via electro-spray ionization, where full-scan mass spectra were collected in positive ion mode from 100-2250 m/z with a spectra rate of 1.0 Hz, 500 V end plate offset, 4200 V capillary, 1.7 bar nebulizer, 8 L/min dry gas, and 180° C dry temp. We also ran tandem mass spectrometry (MS/MS) experiments for each sample with the same full MS settings as described above with the exception of the spectra rate of 1.20 Hz and the 200° C dry temp. We used Bruker Compass Hystar 5.1 software to acquire the data.

### HDX/MS data analysis

We used Compass DataAnalysis 5.1 (Bruker) to identify and deconvolute peptides. For peptide matching and disambiguation, we used PIGEON, using 30 ppm initial cutoff, 7ppm narrow cutoff, 0.020 Da error for MSMS, score cutoff 0.05, peptide length 3-15, charge cutoff 3, 0.1D/e and 0.5min degeneracy cutoffs, using the WT MSMS data. The resultant peptide pool was imported to HDExaminer 3.3.0 (Sierra Analytics, Inc. 2020) and all spectra were fit for each replicate, all peptide assignments and fits were manually checked for quality. Peptide pool results and spectra were exported from HDExaminer and used for peptide-level analysis of both replicates in FEATHER, with envelope centroids from HDExaminer used to produce uptake plots. Deuterated envelope centroids were compared to un-deuterated and to the number of exchangeable residues in each peptide to determine deuteration levels as % of maximum, determined by averaging full-D centroid values across all replicates and mutants for each peptide, or using the theoretical maximum if no full-D spectra were obtained. Deuteration levels of all peptides covering a residue were averaged and subtracted between WT and both mutants and averaged across all timepoints to calculate pseudo-residue level 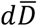.

### HDX/MS experiments summary

| Dataset | SHP2 Rep1 | SHP2 Rep2 | SHP2 Rep3 |
| --- | --- | --- | --- |
| Protein states | WT/Y62D | WT/Y542D,Y580D | Y62D/Y542D,Y580D |
| Date | 10/02/2023 | 02/23/2024 | 01/14/2025 |
| Protease column | Fungal protease XIII/pepsin | Fungal protease XIII/pepsin | ANPEP/pepsin |
| Reaction details | 25mM Tris-HCl, 150mM NaCl, 2mM DTT, temperature 15 °C, final D<sub>2</sub>O concentration 90% |  |  |
| Time course (s) | 0, 30, 120, 480, 960, 1920, 3840, 7680 | 0, 30, 120, 480, 960, 1920, 3840, 7680 | 0, 30, 100, 480, 1920, 7680, 40000 |
| # of time points | 7 | 7 | 6 |
| # of peptides | 213 | 333 | 267 |
| Sequence coverage | 78.9% | 87.9% | 90.5% |
| Average peptide length / Redundancy | 10.3 length, 3.6 redundancy | 12.6 length, 6.9 redundancy | 10.2 length, 4.5 redundancy |

#### • Generation of SHP2 mutant cell lines

The endogenous SHP2 protein was knocked out using CRISPR/Cas9 gRNA from GeneScript (Supplement Table 2). The three sgRNAs were cloned into pLENTICRISPR V2 backbone (RRID:Addgene_140206).

#### • Cloning the sgRNA plasmids

The cloned plasmids control, guide 1,2,3 was transformed into competent *E. coli* DH5⍺ cells (NEB, Cat# C2987H), using the heat shock method. 1uL of DNA was added to 50uL cell aliquot and equilibrated for 5 min. the cells were heat shocked by putting the tube in a water bath set at 42°C for 40 sec. Then, it was placed on ice for 2min. 500uL of SOC media was added and incubated on the shaker at 37°C for an hour. After an hour, the tubes were spun down and 400uL of supernatant was removed. The bacterial pellet was resuspended, and the suspension was added to the antibiotic agar plate. The colonies were picked next day and bacterial DNA miniprep was setup.

#### • Generating lentivirus

Day 0: 293T LentiX cells was seeded in 10cm plate (5-8 million cells/plate) in 10mL DMEM.

Day 1: The amount of plasmid DNA was calculated, JetprimeⓇ transfection media, and packaging plasmids (psPAX2 and pMD2.G) required for transfection (the amount was calculated as per the

protocol given by the manufacturer). The manufacturer’s protocol was followed. Refresh media after 4hrs.

Day 3: The lentivirus was harvested. The media of HEK293T LentiX cell plates was aspirated with a 10mL syringe, and the 0.45µM filter was attached to the syringe, and the virus was filtered out into the falcon tube. (If not using the virus immediately for infection, make aliquots in cryovials and store them in a -80°C freezer).

#### • Infecting the target cells to knock out SHP2

The target cells were seeded (U-2 OS and BT20) in 6 well plates (3 wells for virus + 1 well for no virus control).

Next day, lentivirus infection was performed. The virus was thawed and 1.5mL of the respective virus was added to each well and 1.5mL media to no virus well. 8ug/mL of polybrene was added to each well to increase infection efficiency. The volume was made up to 2mL, and the plates were incubated overnight. The next day, the media was refreshed to remove the virus.

On the following day, the cells were trypsinized if required, and antibiotic selection (blasticidin) was started. The antibiotic selection was performed untill all the cells in the no-virus well were dead.

### Generating isogenic SHP2 mutant cell lines

We performed Gibson cloning to incorporate gRNAs of different SHP2 mutants into the pLX304 vector backbone (Supplementary protocol). The SHP2 mutant cell line generation was done using the same protocol as above, using the JetprimeⓇ transfection and antibiotic selection (puromycin).

## Data availability

Phosphositeplus data is available on https://www.phosphosite.org/homeAction.action. CPTAC data is available at https://proteomic.datacommons.cancer.gov/pdc/. Crystal structure of SHP2 Y62D is deposited on PDB entry 9R16 (https://doi.org/10.2210/pdb9R16/pdb).

## Supporting information

Supplementay protocol

## Acknowledgement

Authors thank Dr. Iok In Christine Chio, Columbia University for providing us with the immortalized MEFs. Authors also thank Dr. Neel Shah for constructive feedback on the manuscript, and Dr. Benjamin Neel, Dr. Jared Johnson, Dr. Hanna Karvonen, and Dr. Shalinda Fernando for helpful discussions.

## Author’s contribution

- Conceptualization: P.K., R.S., A.G., N.V.
- Data curation: P.K., R.S., I.S.G., A.G., N.V.
- Formal Analysis: P.K., R.S., R.R., M.W., S.A., L.C.T., I.S.G., L.C., A.K., W.S.
- Funding acquisition: A.G., N.V.
- Investigation: P.K., R.S., M.W., R.R., I.S.G., W.S., P.H., J.L., A.G., N.V.
- Methodology: P.K., R.S., M.W., R.R., S.A., L.C.T., I.S.G., L.C., A.K., W.S.
- Project administration: A.G., N.V.
- Resources: W.S., P.H., J.L.
- Software: W.S., P.H., A.G., N.V.
- Supervision: N.V.
- Validation: P.K., R.S., L.C.T., I.S.G., P.H., A.G., N.V.
- Visualization: P.K., R.S., M.W., R.R., L.C.T., I.S.G.
- Writing – original draft: P.K., N.V.
- Writing – review & editing: P.K., R.S., M.W., L.C.T., I.S.G., A.G., N.V.

