## Supplementary tables and figures for "A Hotspot Phosphorylation Site on SHP2 Drives Oncoprotein Activation and Drug Resistance"

**Supplementary Table 1:** Crystallographic data collection and refinement statistics

| | SHP2 |
|---|---|
| PDB entry | 9R16 |
| Data processing | |

| | |
|---|---|
| Space group | $P2_1$ |
| Unit cell dimensions<br>a, b, c / Å<br>$\alpha, \beta, \gamma$ / ° | <br>45.17, 212.16, 54.92<br>90.0 96.5 90.0 |
| Resolution[a] | 34.3-2.63 (3.04-2.63) |
| Number of reflections<br>Total<br>Unique | <br>144690 (14234)<br>21059 (2106) |
| $R_{meas}$ | 0.266 (0.834) |
| $R_{pim}$ | 0.101 (0.317) |
| Mean $I/\sigma I$ | 5.7 (2.5) |
| $CC_{1/2}$ | 0.987 (0.718) |
| Multiplicity | 6.9 (6.8) |
| Completeness<br>Spherical<br>Ellipsoidal | <br>69.1 (19.5)<br>89.0 (51.1) |
| Wilson B-factor / Å$^2$ | 54.6 |
| *Refinement* | |
| Resolution | 27.8-2.63 (2.82-2.63) |
| $R_{work}$ | 0.280 (0.305) |
| $R_{free}$ | 0.304 (0.333) |
| Number of atoms | 8499 |
| Average B-factor | 51.0 |
| R.M.S. deviations<br>Bond lengths / Å<br>Bond angles / ° | <br>0.004<br>0.62 |
| Ramachandran plot / %<br>Favored<br>Allowed<br>Outlier | <br>97.6<br>2.2<br>0.2 |
| Clashscore | 0.49 |

**Supplementary Table 2:** gRNA and Primer seuences used in this study

| | Sequences |
|---|---|
| PTPN11_gRNA1 | GATTACTATGACCTGTATGG |
| PTPN11_gRNA2 | GCGCACTGGTGATGACAAAG |
| PTPN11_gRNA3 | TTACTATGACCTGTATGGAG |
| Y62D_Forward | CACTGGTGATgatTATGACCTGTATG |
| Y62D_Reverse | TTCTGAATCTTGATGTGG |
| Y62F_Forward | CACTGGTGATttcTATGACCTGTATG |
| Y62F_Reverse | TTCTGAATCTTGATGTGGG |
| Y542D_Forward | AGGGCACGAAgatACAAATATTAAG |
| Y542D_Reverse | TTCTGAATCTTGATGTGGG |
| Y542F_Forward | AGGGCACGAAttcACAAATATTAAG |
| Y542F_Reverse | TTCCTCTTGCTTTTCTGC |
| Y580D_Forward | TGCTAGAGTCgatGAAAACGTGG |
| Y580D_Reverse | CTGTCTTCTCTCATTTCTGC |
| Y580F_Forward | TGCTAGAGTCttcGAAAACGTGG |
| Y580F_Reverse | CTGTCTTCTCTCATTTCTG |
| pLX304_Forward | AACAGCAGAAAAGTTTCAGAGGTAAGCCTATCCCTAACCCTCT |
| pLX304_Reverse | AACCATCTCCGCGATGTCATTGATCCCGACAGTTAGCCAG |
| SHP2_Ins_pHAT_Forward | tcatcaccatcaccatcacaacactagtagcgctaccatgATGACATCGCGGAGATGGTT |
| SHP2_Ins_pHAT_Reverse | gatttaggtgacactatagaatactcaagcttatgcatgcTCATCTGAAACTTTTCTGCTGTTGC |
| SHP2_Ins_pLX304_Forward | CTGGCTAACTGTCGGGATCAATGACATCGCGGAGATGGTT |
| SHP2_Ins_pLX304_Reverse | CTGGCTAACTGTCGGGATCAATGACATCGCGGAGATGGTT |
| pGEX_sequencing_Forward | TGGTAGAACGAAGCGGCG |
| pGEX_sequencing_Reverse | CGACACCACCACGCTGG |
| His-tag_Forward | caccaccacGGAATTCCGGGCGGGAGG |
| His-tag_Reverse | atgatgatgCATGAATACTGTTTCCTGTGTGAAATTGTTATCC |

**Supplementary Figures:**

Supplementary Figure 1:

**A**

| **Conservation of SHP2 Y62 across animal species** | | | |
|---|---|---|---|
| **Species** | **Common name** | **Sequence** | **Bolded residue** |
| *H. sapiens* | *Human* | KIQNTGD**Y**YDLYGGE | Y62 |
| *P. troglodytes* | *Chimpanzee* | KIQNTGD**Y**YDLYGGE | Y61 |
| *M. musculus* | *House mouse* | KIQNTGD**Y**YDLYGGE | Y62 |
| *M. domestica* | *Housefly* | KIQNTGD**Y**YDLYGGE | Y62 |
| *G. gallus* | *Redfowl* | KIQNTGD**Y**YDLYGGE | Y62 |
| *A. carolinensis* | *Green Anole* | MIRCQDM**K**YDVGGGE | K62 |
| *X. tropicalis* | *Western Clawed Frog* | KIQNTGD**Y**YDLYGGE | Y62 |
| *D. rerio* | *Zebrafish* | KIQNTGD**Y**YDLYGGE | Y62 |
| *D. melanogaster* | *Fruitfly* | KIQNNGD**F**FDLYGGE | F62 |
| **SHP1** | | | |
| *H. sapiens* | | RIQNSGD**F**YDLYGGE | F60 |

**B**

| **Conservation of Y62 in nSH2 domain in *Homo sapiens*** | | | |
|---|---|---|---|
| **Proteins** | **Sequence** | **Phosphosite** | **Frequency** |
| SHP2 | KIQNTGD**Y**YDLYGGE | Y62 | 2116 |
| RASA1 | IIAMCGD**Y**YIGGRRF | Y239 | 3 |
| YES | RKLDNGG**Y**YITTRAQ | Y222 | 1816 |
| FYN | RKLDNGG**Y**YITTRAQ | Y213 | 1819 |
| FGR | RKLDMGG**Y**YITTRVQ | Y208 | 306 |
| LYN | RSLDNGG**Y**YISPRIT | Y193 | 787 |
| BLK | RCLDEGG**Y**YISPRIT | Y187 | 148 |
| SLAP/SLA | FRLPNNW**Y**YISPRLT | Y142 | None reported |
| SH2D1B | IFREKHG**Y**YRIQNSN | Y62 | None reported |
| ITK | TNDNPKR**Y**YVAEKYV | Y305 | None reported |
| BTK | CSTPQSQ**Y**YLAEKHL | Y344 | 189 |
| TEC | TTTSPKK**Y**YLAEKHA | Y312 | None reported |

**(A)** Sequence alignments of SHP2 orthologs across species, and human SHP1. (**B**) Sequence alignments of human SH2 domains with phosphosite and frequency indicated.

Supplementary Figure 2:

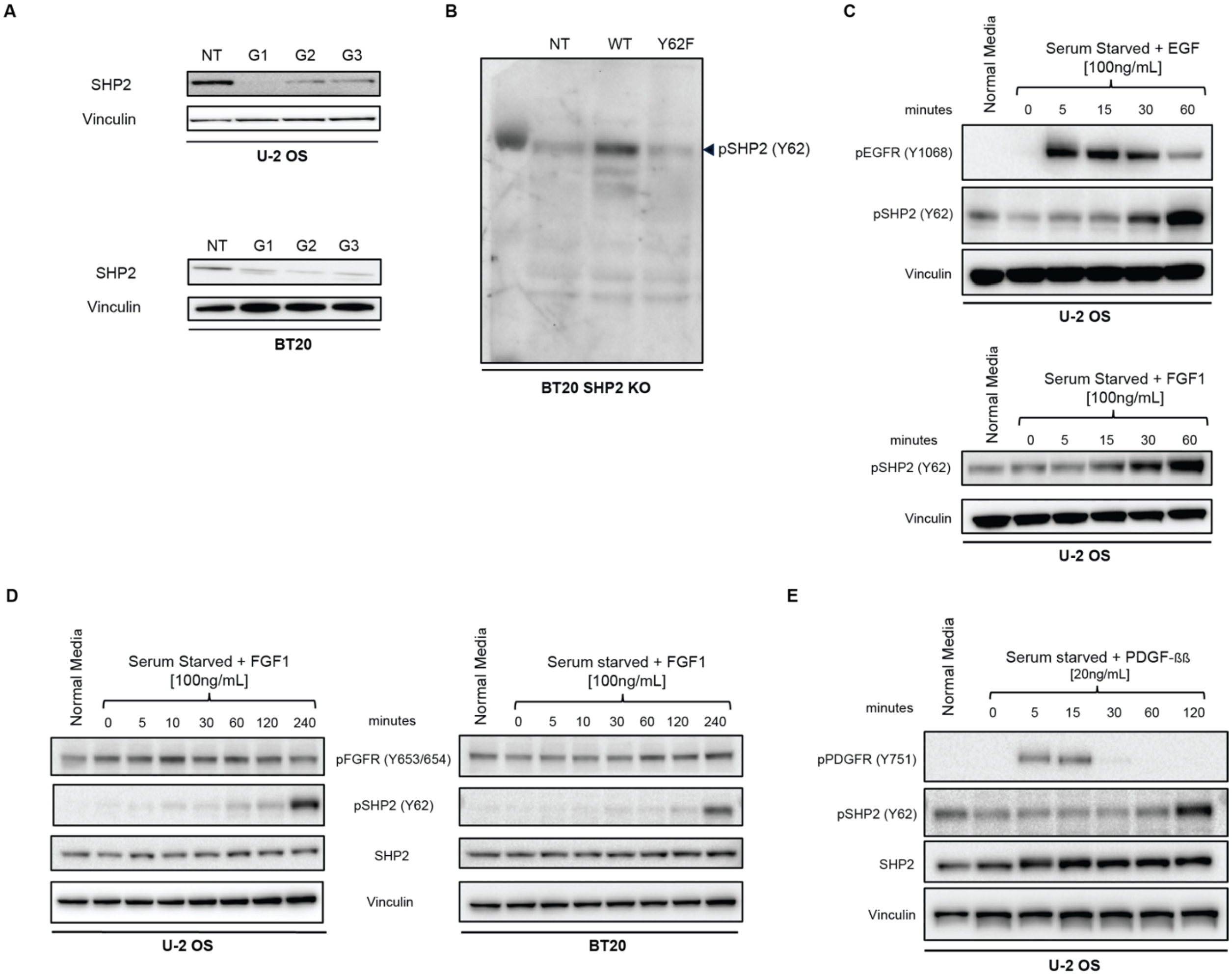

**(A)** Immunoblot analysis of BT20 and U2-OS cells bearing SHP2 knock out. NT, non-targeting guide RNA. (**B)** Immunoblot analysis of pSHP2 Y62 antibody in BT20 SHP2 knock out cells overexpressing SHP2 WT and Y62F. (**C-E**) Immunoblot analysis of indicated cell lines serum starved (24 hours) then treated with (**C**) EGF **(**100 ng/mL), (**D**) FGF1 (100ng/mL), and (**E**) PDGF-ββ (20ng/mL) for the indicated timepoints.

Supplementary Figure 3:

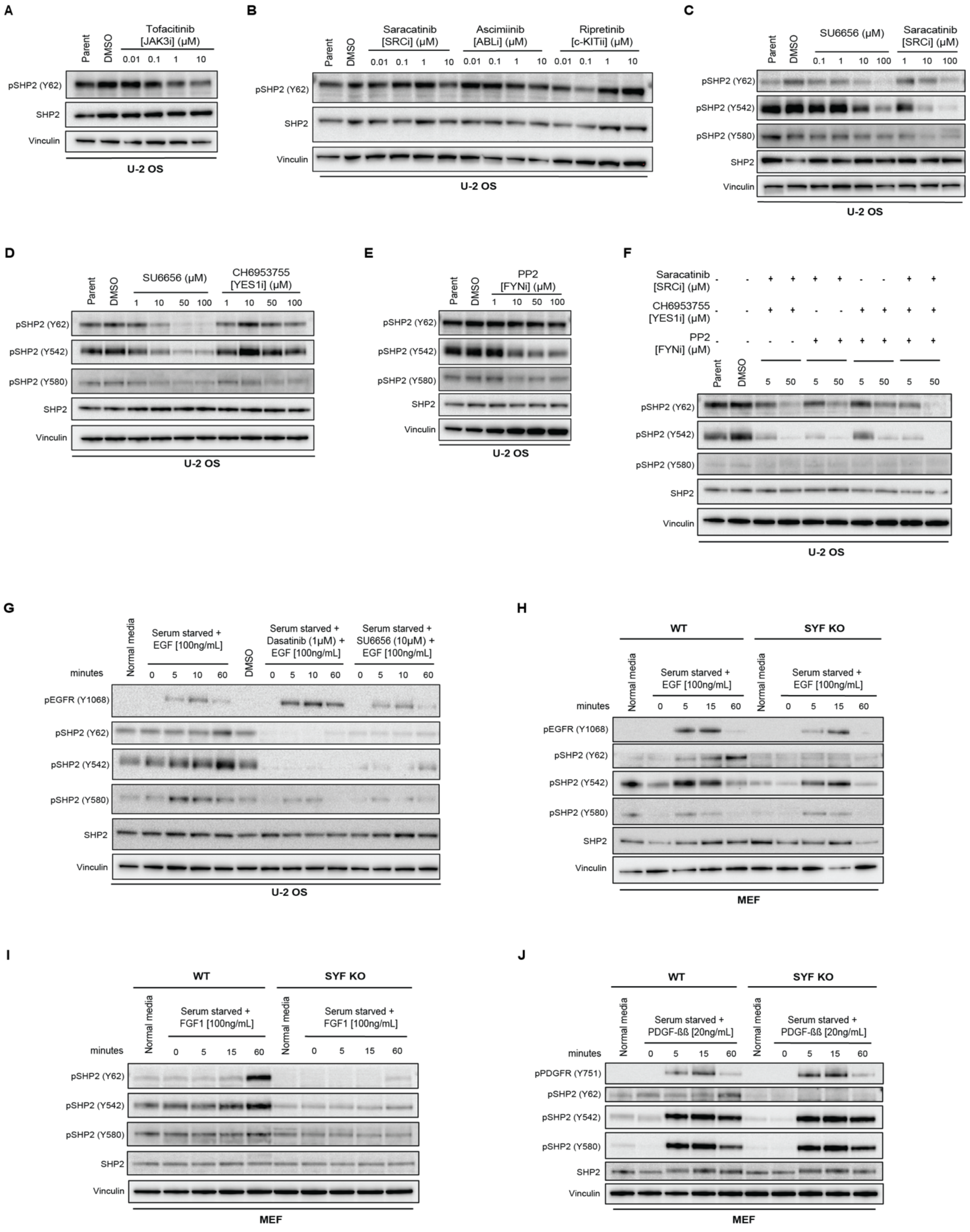

(**A-F**) Immunoblot analysis of U-2 OS cells treated with (**A**) tofacitinb, (**B**) saracatinib, asciminib, and ripretinib, (**C**) SU6656 and saracatinib, (**D**) SU6656 and CH6953755, (**E**) PP2, (**F**) and double and triple combinations of saracatinib, CH6953755 and PP2 at indicated concentrations, and DMSO. (**G**) Immunoblot analysis of U-2 OS cells, serum-starved (24 hours) and treated with dasatinib and SU6656, followed by EGF (100 ng/mL) for indicated timepoints. (**H-J**) Immunoblot analysis of SYF knock out and wildtype MEFs, serum-starved and treated with (**H**) EGF (100 ng/mL), (**I**) FGF1 (100 ng/mL), and (**J**) PDGF-ββ (20 ng/mL).

Supplementary Figure 4:

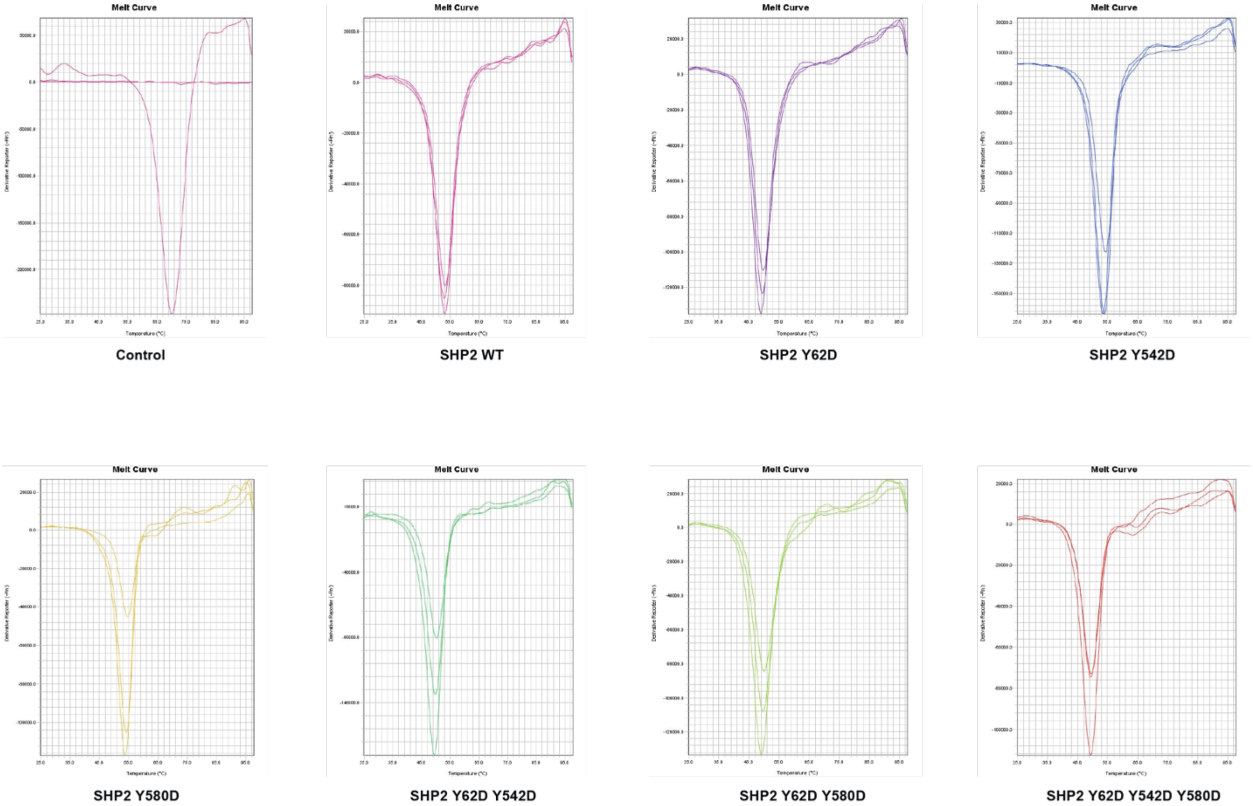

Melting temperatures ($T_m$) of wild-type and mutant SHP2 proteins by differential scanning fluorimetry (DSF) (n = 3).

Supplementary Figure 5:

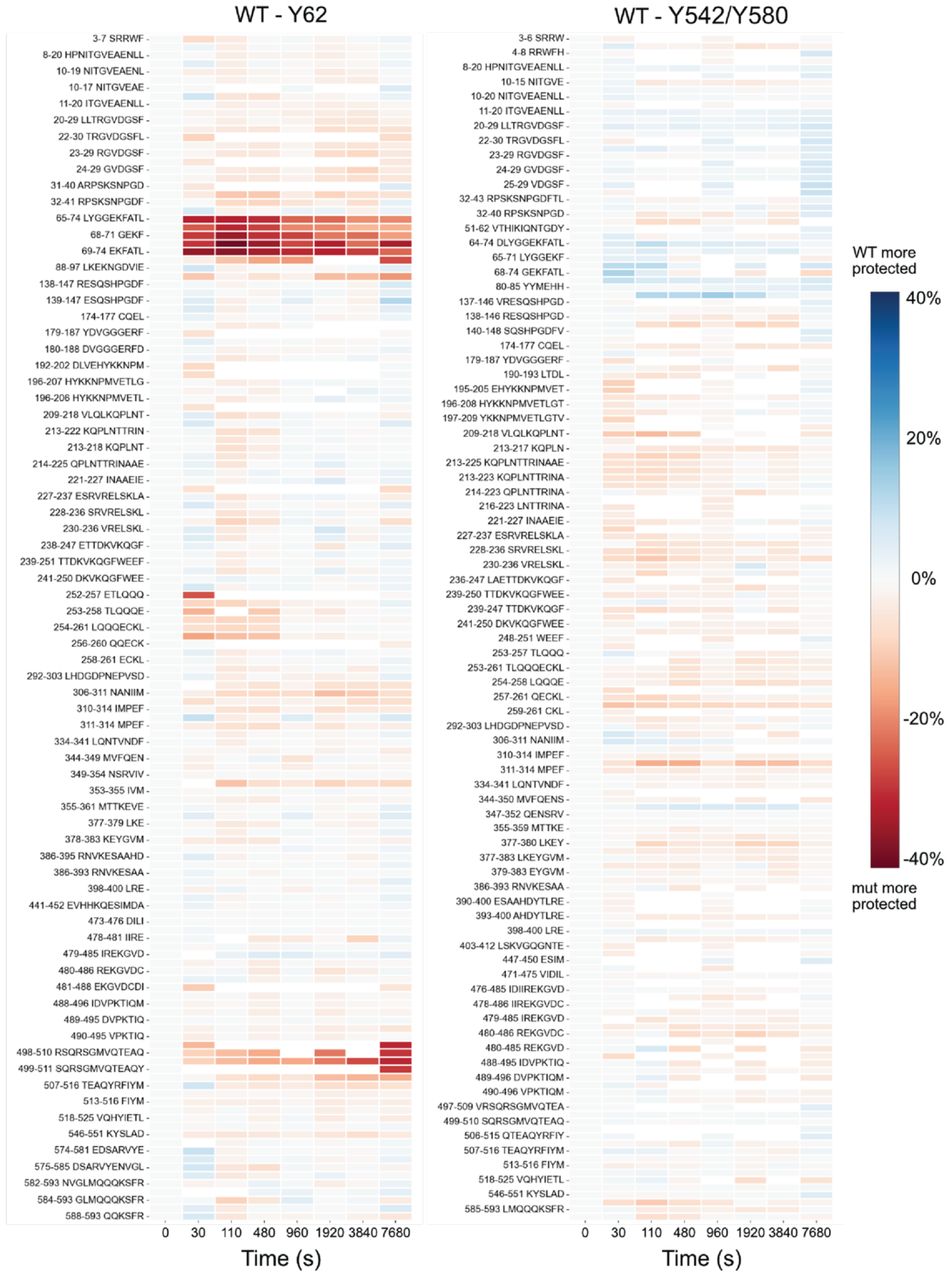

Peptide-level exchange vs time-course heat plots comparing SHP2 WT to the two SHP2 mutants, Y62D and Y542D/Y580D. Plots show peptide length-normalized exchange at each timepoint.
