## Supplementay protocol for "A Hotspot Phosphorylation Site on SHP2 Drives Oncoprotein Activation and Drug Resistance"

Protocol – Gibson cloning of SHP2 mutants into pLX304

### Overview

The protocol consists of 3 PCR steps and 1 Gibson reaction

### PCR 1: Amplifies part 1 of the pLX304 backbone

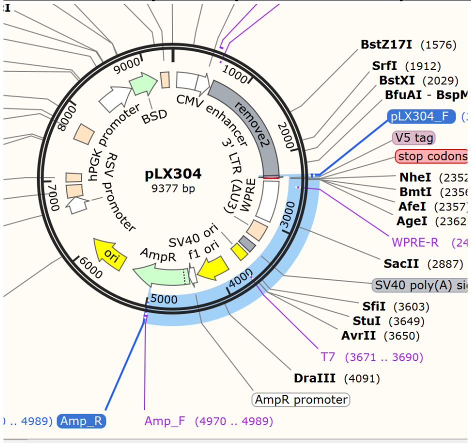


### PCR 2: Amplifies part 2 of the pLX304 backbone

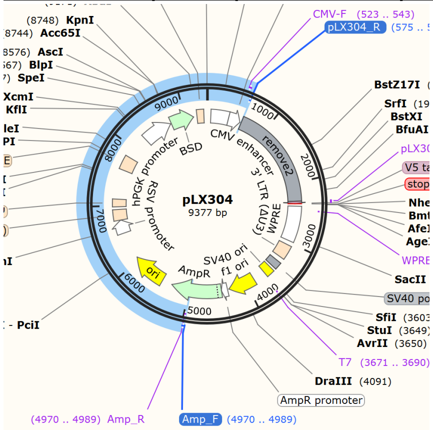


### PCR 3: Amplifies the SHP2 variants of the pGEX vectors

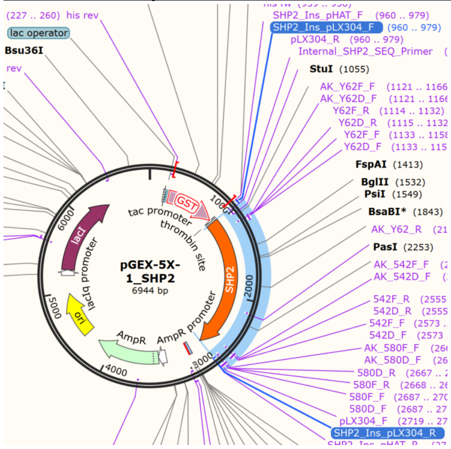

**Materials**
Gibson 2x MM
Primers
- Amp_F
- Amp_R
- pLX304_F
- pLX304_R
- SHP2_Ins_pLX304_F
- SHP2_Ins_pLX304_R

Plasmids
- pLX304
- pGEX_SHP2_WT
- pGEX_SHP2_Y62F
- pGEX_SHP2_Y542F
- pGEX_SHP2_Y580F
- pGEX_SHP2_Y62F_Y542F
- pGEX_SHP2_Y62F_Y580F
- pGEX_SHP2_Y542F_Y580F
- pGEX_SHP2_Y62F_Y542F_Y580F

1% Agarose gel
DpnI
PCR clean up kit
Gibson 2x HIFI MM
*E. coli* Dh5alpha

**Methods**
PCR 1 – 200 µL – **2690 bp** – 68 degrees annealing

Prepare in a PCR tube:

97 uL Nuclease Free Water
1 uL pLX304**_F** (100 µM)
1 uL Amp**_R** (100 µM)
1 uL pLX304 S (~200 ng/µL)
100 uL Q5 2x MM

Vortex – spin down – Divide 100 µL over 2 PCR tubes

PCR: 30 cycles of step 2-4
98 30 seconds
98 10 seconds
68 15 seconds
72 **1.5** minutes
72 2 minutes
10 hold

PCR 2 – 200 uL – **5000 bp** – 68 degrees annealing

Prepare in a PCR tube:

97 µL Nuclease Free Water
1 µL pLX304**_R** (100 µM)
1 µL Amp**_F** (100 µM)
1 µL pLX304 S (~200 ng/µL)
100 µL Q5 2x MM

*Really watch that you are adding the 1 µL!

Vortex – spin down – Devide 100 µL over 2 PCR tubes

PCR: 30 cycles of step 2-4
98 30 seconds
98 10 seconds
68 15 seconds
72 **5** minutes
72 **10** minutes
10 hold
PCR 3 – 8x 100 µL – **1783 bp** – 68 degrees annealing
Prepare MM in a 1.5 mL tube:

388 µL Nuclease Free Water

4 µL SHP2_Ins_pLX304_**F** (100 µM)
4 µL SHP2_Ins_pLX304_**R** (100 µM)
400 µL Q5 2x MM

Vortex – spin down - Devide over 8 PCR tubes: 98 µL each (just to make sure you don't have to little for the last one)

Add 0.5 µL of each pGEX SHP2 plasmid to a tube – watch closely.

pGEX_SHP2_WT = 1
pGEX_SHP2_Y62F = 2
pGEX_SHP2_Y542F = 3
pGEX_SHP2_Y580F = 4
pGEX_SHP2_Y62F_Y542F = 5
pGEX_SHP2_Y62F_Y580F = 6
pGEX_SHP2_Y542F_Y580F = 7
pGEX_SHP2_Y62F_Y542F_Y580F = 8

Vortex – spin down

PCR: 30 cycles of step 2-4
98 30 seconds
98 10 seconds
**67** 15 seconds
72 **1** minutes
72 2 minutes
10 hold

---

<u>1% Agarose gel control</u>
Take 5 µL of each unique sample (so for PCR 1 and PCR2 only 1 check, for PCR 3 all 8 samples) to a gel – check if the size is roughly correct.

5 µL PCR
5 µL H2O
2 µL 6x Loading dye

Left over 95 µL PCR can be stored at 4 degrees if not immediately continuing to DpnI treatment.

<u>DpnI digestion</u>

To each PCR tube, add 1 µL DpnI
Incubate at 37 degrees in the PCR machine for 30 minutes

<u>PCR clean up</u>

Immediately following DpnI digest – clean up the PCR/DpnI samples

Use Nucleaspin Gel and PCR clean up

*For PCR 1/2*
Add 400 µL yellow binding buffer to a 1.5 mL tubes
Combine and add 200 µL PCR reaction

*For PCR 3*
Add 200 µL yellow binding buffer to a 1.5 mL tubes (1 for each PCR tube)
Add 100 µL PCR reaction

Follow protocol. Elute with nuclease free water. Elute 20 µL for PCR 1 and 2, 15 µL for each PCR 3.

<u>Measure concentrations – can be stored at -20.</u>

<u>Gibson</u>

*Volumes depend on concentrations of DNA – lets decide when you know.

In PCR tube

1 µL Backbone PCR1
2 µL Backbone PCR2
1 µL SHP2 Insert DNA
5 µL 2x HIFI MM

Incubate for 1 hour at 50 degrees Celsius (lid at 80 degrees).

Gibson can be stored at -20 if not immediately continuing to transformation.

<u>Transformation</u>
Transform 2.5 µL to 50 µL dh5x cells

As per protocol (30 seconds at 42 heatshock)

Plate all cells to ampicillin plates.
